# Control of cytokinesis by β-adrenergic receptors indicates an approach for regulating cardiomyocyte endowment

**DOI:** 10.1101/782920

**Authors:** Honghai Liu, Cheng-Hai Zhang, Niyatie Ammanamanchi, Sangita Suresh, Christopher Lewarchik, Krithika Rao, Gerrida M. Uys, Lu Han, Maryline Abrial, Dean Yimlamai, Balakrishnan Ganapathy, Christelle Guillermier, Nathalie Chen, Mugdha Khaladkar, Jennifer Spaethling, James H. Eberwine, Junhyong Kim, Stuart Walsh, Sangita Choudhury, Kathryn Little, Kimberly Francis, Mahesh Sharma, Melita Viegas, Abha Bais, Dennis Kostka, Jun Ding, Ziv Bar-Joseph, Yijen Wu, Matthew L. Steinhauser, Bernhard Kühn

## Abstract

One million patients with congenital heart disease (CHD) live in the US. They have a lifelong risk of developing heart failure. Current concepts do not sufficiently address mechanisms of heart failure development specifically for these patients. We show that cardiomyocyte cytokinesis failure is increased in tetralogy of Fallot with pulmonary stenosis (ToF/PS), a common form of CHD. Labeling of a ToF/PS baby with isotope-tagged thymidine showed cytokinesis failure after birth. We used single-cell transcriptional profiling to discover that the underlying mechanism is repression of the cytokinesis gene ECT2, and show that this is downstream of β-adrenergic receptors (β-AR). Inactivation of the β-AR genes and administration of the β-blocker propranolol increased cardiomyocyte division in neonatal mice, which increased the endowment and conferred benefit after myocardial infarction in adults. Propranolol enabled the division of ToF/PS cardiomyocytes. These results suggest that β-blockers should be evaluated for increasing cardiomyocyte division in patients with ToF/PS and other types of CHD.

## INTRODUCTION

Congenital heart disease (CHD) is the most common birth defect. Improvements in diagnosis and treatment of CHD have increased survival, and 1 million patients live in the US with CHD^1^. Patients with CHD have a high lifetime risk for developing heart failure, and current thinking about this is based on research on heart failure in adult patients^1^. We have considered the possibility that CHD may alter cellular growth of the myocardium, i.e., cardiomyocyte proliferation and differentiation, because these mechanisms are active in infants and children without heart disease^2, 3^.

Tetralogy of Fallot with pulmonary stenosis (ToF/PS) is a common form of CHD with relatively uniform structural defects (anterior deviation of the infundibulum, pulmonary stenosis, ventricular septal defect, and right ventricular hypertrophy). Despite extensive progress in understanding the genetic causes of CHD, the majority of ToF/PS remains genetically unexplained^4^. Infants and children with ToF/PS rarely have heart failure, but it is a well-documented cause of morbidity and mortality in adults^5–12^. Current thinking is that the sequelae of cardiac surgery cause an increased risk of heart failure. However, classical studies showed severe cardiomyocyte changes in ToF/PS patients prior to surgery^13, 14^, but did not examine cardiomyocyte proliferation and differentiation. This suggested to us that myocardial changes happen before surgery. We have considered this possibility and found changes in cardiomyocyte proliferation and differentiation in ToF/PS.

Recent studies have shown that cardiomyocytes divide in human infants and children in contrast to the extremely low cardiomyocyte division rate in adults^2, 3, 15^. When cardiomyocytes stop proliferating, they undergo incomplete cell cycles, leading to binucleated cardiomyocytes. Although the mechanisms of formation of binucleated cardiomyocytes are unknown, it is thought that they do not divide further^16^. Mice and rats form binucleated cardiomyocytes in the first week after birth^17–19^. Zebrafish have only mononucleated cardiomyocytes, which can divide and regenerate myocardium^20^, which has led to the hypothesis that a high percentage of mononucleated cardiomyocytes is the foundation for myocardial regeneration. However, humans have 70% mononucleated cardiomyocytes and yet do not regenerate myocardium. In addition, cardiomyocytes differentiate by endocycling, which increases the DNA content of nuclei without nuclear division, i.e., they become polyploid. Humans show a high degree of endocycling around 10 years after birth^2, 3, 15^. Two recent papers have altered the percentage of binucleated cardiomyocytes in mice and zebrafish, but this resulted in additional large changes of polyploid cardiomyocytes^21, 22^. Although multiple studies have suggested that cardiomyocytes become binucleated by incomplete cytokinesis^23–25^, the precise mechanisms and relevance of cardiomyocyte binucleation are still unknown. During cytokinesis, a contractile ring forms at the future division plane^26^. Contraction of this ring is triggered by the cytokinesis protein ECT2, a RhoA guanine-nucleotide exchange factor. RhoA-GTP activates, via Rho-associated protein kinase (ROCK), non-muscle myosin II, which constricts the cleavage furrow.

Different pathways regulate cardiomyocyte proliferation, with the Hippo tumor suppressor pathway taking a central position^27^. The Hippo pathway is regulated by G protein-coupled receptors (GPCR), and, in the heart, activated by β-adrenergic receptors (β-AR)^28^. β-AR regulate cardiomyocyte contractile function, by adjusting the intracellular second messenger cyclic adenosine monophosphate (cAMP)^29^. In CHD, and specifically in ToF/PS, β-AR signaling is overactivated^30–34^. Adrenergic signaling has also been evaluated in the context of heart regeneration in mice^35, 36^ and cell cycle activity in cultured rat cardiomyocytes^37–39^; however, these results were obtained without genetic disruption of signaling pathways. Our results extend these findings by demonstrating a function of β-adrenergic receptors (β-AR) signaling in regulating cardiomyocyte cytokinesis *in vivo*.

Using formation of binucleated cardiomyocytes as read-out for the definitive endpoint of cell division, we discovered extensive changes of cardiomyocyte proliferation in ToF/PS. We identified the mechanisms of formation of binucleated cardiomyocytes, establishing a new connection between β-AR signaling and regulation of cardiomyocyte cytokinesis.

## RESULTS

### Infants with Tetralogy of Fallot with pulmonary stenosis have increased binucleated cardiomyocytes

We examined samples from the right ventricle of patients with ToF/PS and made the surprising observation that the percentage of binucleated cardiomyocytes was increased to 50-60% (**Fig. 1A-C**), suggesting extensively increased cytokinesis failure. The temporal pattern of this increase shows that babies with ToF/PS were born with the appropriate percentage of 20% binucleated cardiomyocytes^2, 3^, but that the increase happened in the first 6 months after birth (**Fig. 1B**). All ToF/PS patients > 2 months had cardiomyocytes with > 2 nuclei, a very rare phenotype in humans without heart disease (**Fig. 1C**), suggesting that multiple serial cytokinesis failures occurred. Bi- and multi-nucleated cardiomyocytes were present in 6 and 13-year-old patients, i.e., after the decline of cardiomyocyte cell cycle activity to the very low levels present in adults. This shows that bi- and multi-nucleated cardiomyocytes generated in the first 6 months after birth live for at least one decade. To directly assess the generation of mono- and binucleated cardiomyocytes, we labeled a 1-month-old ToF/PS baby with ^15^N-thymidine and examined uptake and retention with multiple-isotope imaging mass spectrometry (MIMS) at 7 months of age (**Fig. 1D, E**). ^15^N-thymidine labeling was twice as high in the binucleated cells compared to mononucleated cardiomyocytes (**Fig. 1F**), indicating extensive cardiomyocyte cytokinesis failure corresponding to a 20-30% reduction of the number of cardiomyocytes (endowment). These findings motivated us to determine the mechanisms controlling cytokinesis in cardiomyocytes.

**Figure 1.**
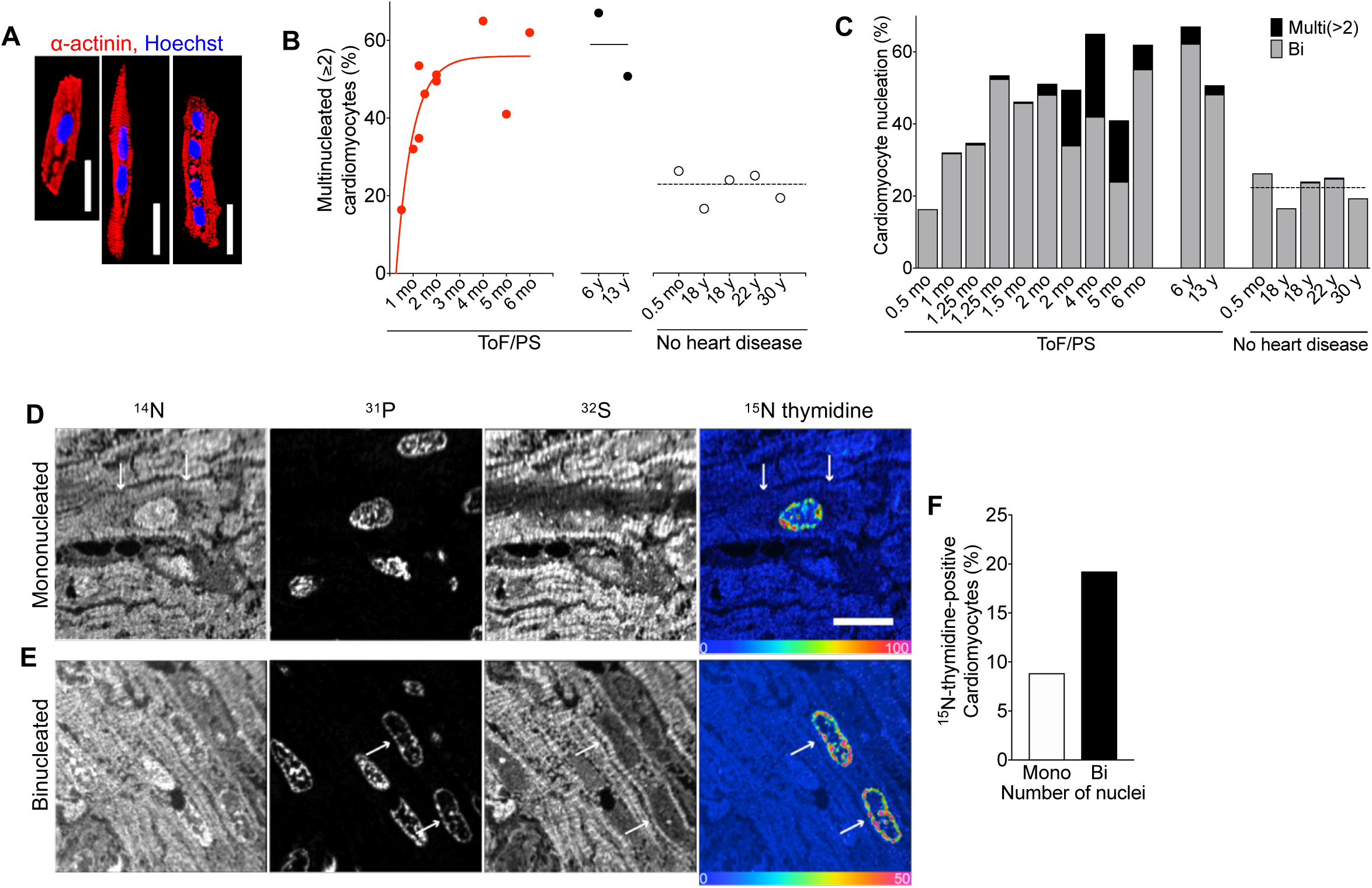
Cardiomyocytes in infants with Tetralogy of Fallot with pulmonary stenosis (ToF/PS) fail to divide. (**A-C**) Patients with Tetralogy of Fallot and pulmonary stenosis (ToF/PS) show an increased proportion of multinucleated (≥2) cardiomyocytes (filled symbols and solid lines in **B**). Each symbol in (**B**) and bar in (**C**) represents one human heart (ToF/PS: n = 12; No heart disease: n = 5). (**D-F**) A 4-week-old infant with ToF/PS was labeled with oral ^15^N-thymidine. Myocardium was analyzed by multiple-isotope imaging mass spectrometry (MIMS) at 7 months. (**D-E**) ^31^P reveals nuclei and ^32^S morphologic detail, including striated sarcomeres. Nuclei are dark in the ^32^S image. The ^15^N/^14^N ratio image reveals ^15^N-thymidine incorporation. The blue end of the scale is set to natural abundance (no label uptake) and the upper bound of the rainbow scale is set to 50% above natural abundance. The white arrows in (**D**) indicate the boundaries of a labelled mononucleated cardiomyocyte and in (**E**) the nuclei of a binucleated cardiomyocyte. (**F**) The percentage of labeled binucleated cardiomyocytes is higher than mononucleated cardiomyocytes (Mononucleated cardiomyocytes analyzed: n = 282; 15N+ mononucleated cardiomyocytes: n = 25; Binucleated cardiomyocytes analyzed: n = 104 (208 total nuclei); 15N+ binucleated cardiomyocytes: n = 20 (40 total nuclei)), indicating that this patient experienced extensive cytokinesis failure. Scale bar: 20 µm (**A**), 10 µm (**D**).

### Cytokinesis failure in cardiomyocytes is associated with low levels of the Rho guanine nucleotide exchange factor Ect2

To determine the cellular mechanisms of cytokinesis failure in cardiomyocytes, we performed live cell imaging with neonatal rat ventricular cardiomyocytes that undergo binucleation (NRVM, **Fig. 2A, Video S1**). Cleavage furrow ingression was observed in 80% of the cardiomyocytes studied, followed by cleavage furrow regression. We used a transgenic mouse model expressing the fluorescent ubiquitination-based cell cycle indicator (FUCCI) to highlight cell cycle progression ^40^, which showed normal cell cycle progression until cleavage furrow regression (**Videos S2, S3**). This finding demonstrates that failure of abscission generates binucleates from mononucleated cardiomyocytes.

**Figure 2.**
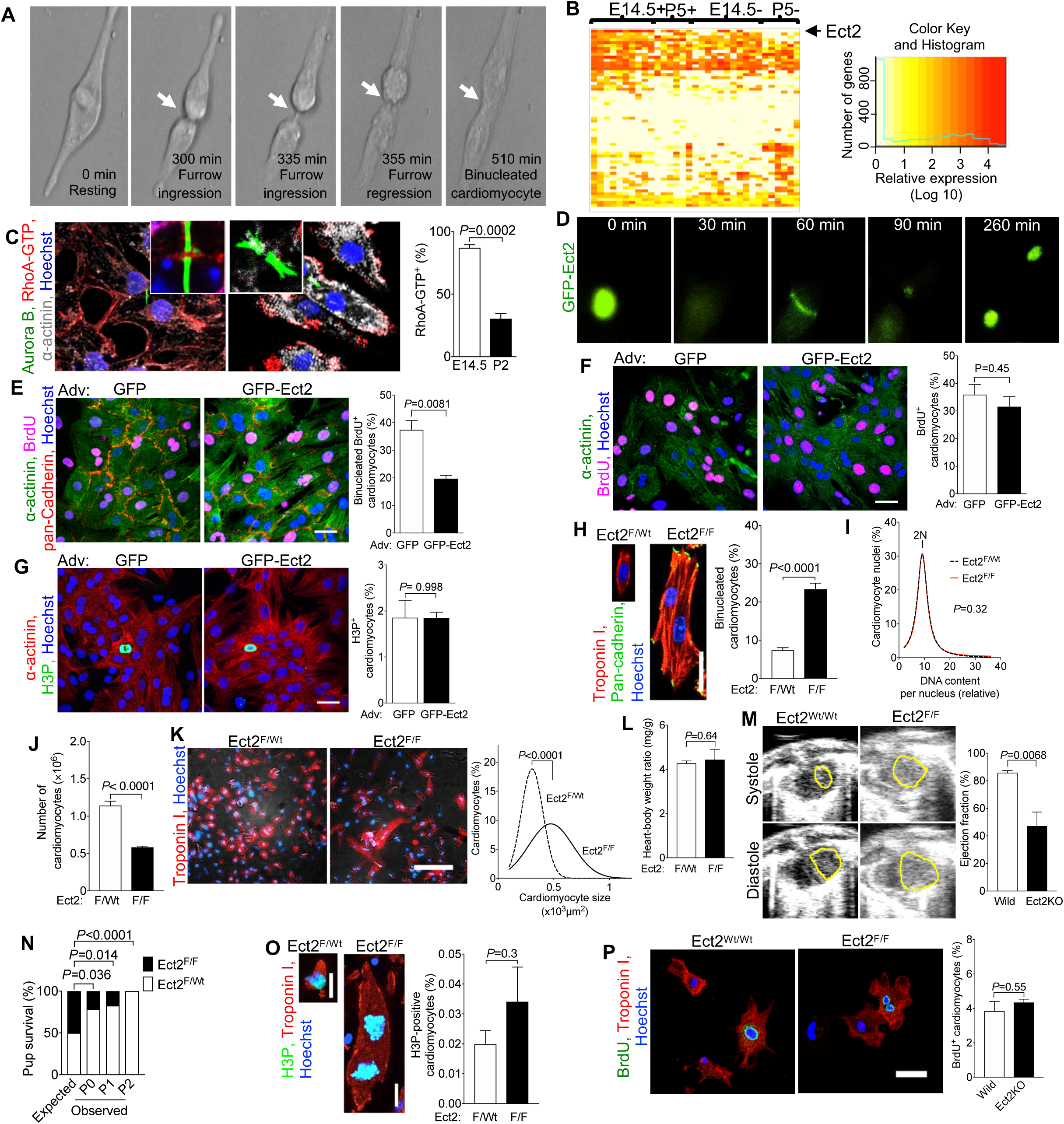
Ect2 levels regulate cardiomyocyte cytokinesis and Ect2 gene inactivation lowers endowment and is lethal in mice. (**A**) Live cell imaging of neonatal rat cardiomyocytes (NRVM, P2-P3, 52 cardiomyocytes) shows that cleavage furrow regression precedes formation of binucleated cardiomyocytes, corresponding to **Video S1**. Cleavage furrow ingression is between 300-335 min, regression at 355 min, and formation of a binucleated cardiomyocyte at 510 min. (**B**) Transcriptional profiling of single cycling (+) and not cycling (-) cardiomyocytes at embryonic day 14.5 (E14.5) and 5 days after birth (P5) reveals that of 61 Dbl-homology family RhoGEF, Ect2 is significantly repressed in cycling P5 cardiomyocytes (P <0.05). (**C**) Binucleating cardiomyocytes exhibit lower RhoA activity (RhoA-GTP) at the cleavage furrow (E15.4: 59 midbodies; P2: n = 56 midbodies). (**D**-**G**) NRVM were transduced with Adv-CMV-GFP-Ect2. Live cell imaging shows appropriate and dynamic localization of GFP-ECT2 in cycling NRVM (**D**, corresponding to **Video S4**), reduced cytokinesis failure and reduced generation of binucleated cardiomyocytes. (**F, G**) Overexpression of GFP-Ect2 does not alter cardiomyocyte S-(**F,** GFP: n = 407; GFP-Ect2: n = 285) or M-phase (**G,** GFP: n = 432; GFP-Ect2: n = 443). (**H**-**P**) Ect2 gene inactivation in the αMHC-Cre; Ect2^F/F^ mice at P1 (**H-O)** showed increased binucleated cardiomyocytes (**H**, Ect2^F/wt^ n = 6, Ect2^F/F^ n = 6 hearts) without change of DNA content per nucleus (**I**, Ect2^F/wt^ n = 642 cardiomyocytes, Ect2^F/F^ n = 647 cardiomyocytes), and a 50% lower cardiomyocyte endowment (**J**, Ect2^F/wt^ n = 12, Ect2^F/F^ n = 5 hearts). The reduced endowment triggers compensatory cardiomyocyte hypertrophy (**K**, Ect2^F/wt^ n = 1,138 cardiomyocytes from 6 hearts, Ect2^F/F^ n = 1,015 cardiomyocytes from 6 hearts), without change of heart weight (**L**, Ect2^F/wt^ n = 14, Ect2^F/F^ n = 6 hearts). (**M, N**) The lower cardiomyocyte endowment leads to myocardial dysfunction at P0 (**M**, left ventricular endocardium outlined in yellow, Ect2^wt/wt^ n = 4, Ect2^F/F^ n = 3 mice) and lethality before P2 (**N, Video S5**), but does not alter the M-phase (**O**, Ect2^F/wt^ n = 6, Ect2^F/F^ n = 6 hearts) and cell cycle entry (**P**, BrdU uptake, Ect2^flox^ gene inactivation with αMHC-MerCreMer, tamoxifen DOL 0, 1, 2, followed by 3 days culture). Statistical significance was tested with Student’s *t*-test if not specified, and Fisher’s exact test (**N**). Scale bars 30 μm (**E, P**), 50 μm (**F, G**), 100 μm (**K**).

To identify the molecular mechanisms of cleavage furrow regression, we separated cycling from non-cycling cardiomyocytes and took a single cell transcriptional profiling approach to compare the expressed genes (**Fig. S1**). We isolated embryonic (Embryonic day 14.5, E14.5) and neonatal (Postnatal day 5, P5) cardiomyocytes and identified cycling cardiomyocytes with the mAG-hGem reporter of the FUCCI indicator ^40^(**Fig. S2**). We performed deep, genome-wide, single-cell transcriptional analysis with the Eberwine method ^41–43^, followed by validation of the results (**Fig. S3**). Because RhoA activation is required for cleavage furrow constriction, we examined the expression of Dbl-homology Rho-Guanine Nucleotide Exchange Factors (GEF) in the single cell transcriptional dataset (**Fig. 2B, Table S1**). Ect2 mRNA was present in cycling E14.5 cardiomyocytes but not in binucleating P5 cardiomyocytes (**Fig. 2B**). Other genes controlling cytokinesis, *i.e.*, *Racgap1* (inactivating RhoA), *RhoA*, *Anillin*, *Aurkb,* and *Mklp1,* were present in P5 cycling cardiomyocytes (**Fig. S4**), indicating that *Ect2* is uniquely regulated. In accordance with the decreased Ect2 levels, active RhoA (RhoA-GTP) was decreased in binucleating cardiomyocytes (**Fig. 2C**). Taken together, these results show insufficient Ect2 levels in cardiomyocytes lead to less RhoA activation, weakening their cleavage furrows^26^.

We tested whether increasing Ect2 expression enables cardiomyocyte abscission by expressing *GFP-Ect2* ^44^. Live cell imaging showed the functionality of GFP-ECT2 in cardiomyocytes (**Fig. 2D** and **Video S4)** and increased cardiomyocyte abscission (**Fig. 2E**) without inducing apoptosis (**Fig. S6**). *GFP-Ect2* did not alter cardiomyocyte BrdU uptake (**Fig. 2F**) or H3P (**Fig. 2G**). In conclusion, increasing *Ect2* expression in cardiomyocytes has a specific effect on abscission without changing cell cycle entry or progression.

### Lowering Ect2 expression reduces cardiomyocyte endowment and heart function

We next tested the hypothesis that lowering the expression of Ect2 induces cytokinesis failure *in vivo*. To this end, we inactivated the Ect2^flox^ gene in mice with αMHC-*Cre* ^45^, (**Fig. S6A**). *αMHC-Cre; Ect2^flox/flox^* mice showed a 3.2-fold increase of binucleated cardiomyocytes (23.3%, **Fig. 2H**), compared to *αMHC-Cre; Ect2^wt/flox^* mice (7.4%, P<0.0001), at P1. *Ect2* inactivation did not change the DNA contents of nuclei (**Fig. 2I**). *αMHC-Cre^+^;Ect2^flox/flox^* pups had 583,000 ± 15,379 cardiomyocytes (n=5 hearts) at P1, a 49% decrease compared to *αMHC-Cre^+^;Ect2^wt/flox^* mice (1,140,833 ± 58,341 cardiomyocytes, n=12 hearts, P<0.0001, **Fig. 2J**). The mean cardiomyocyte size in *αMHC-Cre;Ect2^flox/flox^* mice was increased by 65% (**Fig. 2K**). These results show that the lower endowment in *αMHC-Cre;Ect2^flox/flox^* pups triggered cardiomyocyte hypertrophy, and not a compensatory increase in cell cycling. The heart weight was unchanged (**Fig. 2L**). Echocardiography showed that *αMHC-Cre^+^; Ect2^flox/flox^* had a significantly decreased heart function, measured by ejection fraction (EF=49.6%), compared with control (EF=85.9%, **Fig. 2M**, **Video S6** and **S7**). All the *αMHC-Cre^+^; Ect2^flox/flox^* died before P2 (**Fig. 2N, S6B**). *Ect2^flox^* inactivation did not change cardiomyocyte M-phase activity, as measured by quantification of H3P-positive nuclei (**Fig. 2O**). We observed binucleated Ect2^flox/flox^ cardiomyocytes with both nuclei being in M-phase, indicating that forcing cytokinesis failure does not prevent entry into another cell cycle and advancement to karyokinesis (**Fig. 2O**). This finding suggests a mechanism for how cardiomyocytes with four and more nuclei are generated, *i.e.*, by serial karyokinesis and failure of abscission. To determine whether Ect2 gene inactivation alters cardiomyocyte cell cycle entry, we inactivated an Ect2^flox46^ gene with αMHC-MerCreMer^47^ (tamoxifen P0, 1, 2) *in vivo*, thus circumventing the lethality of inactivating with αMHC-Cre. We isolated cardiomyocytes at P2 and cultured for 3 days in the presence of BrdU. Ect2 inactivation did not alter cell cycle entry (**Fig. 2P**) or cell viability (**Fig. S9**). In conclusion, Ect2 inactivation induced cytokinesis failure in cardiomyocytes, which decreased endowment by 50% and led to severely decreased ejection fraction and death.

### ß-adrenergic receptors control cardiomyocyte abscission and endowment by regulating the Hippo tumor suppressor pathway and *Ect2*

We next sought to identify the mechanisms responsible for decreasing transcription of the *Ect2* gene. Previous publications suggested that the Hippo tumor suppressor pathway regulates cardiomyocyte proliferation ^48–50^. YAP1, the central transcriptional co-regulator controlled by the Hippo pathway, forms a protein complex with TEAD transcription factors ^49^. We identified five binding sites for the transcription factors TEAD 1 and 2 in the *Ect2* promoter (**Fig. S7**). Removing these TEAD-binding sites individually or *en bloc* decreased Ect2 promoter activity in Luciferase assays (**Fig. 3A**). siRNA knockdown of TEAD1/2 reduced Ect2 mRNA levels and increased the proportion of binucleated cardiomyocytes (**Fig. 3B-C**). Adenoviral-mediated overexpression of wild type YAP1 (YAP1-WT) and a non-degradable version (YAP1-S127A) in NRVMs increased Ect2 mRNA levels and reduced the proportion of binucleated cardiomyocytes (**Fig. 3D-E**). These results show that YAP1 and TEAD1/2 regulate the expression of Ect2 and cardiomyocyte abscission.

**Figure 3.**
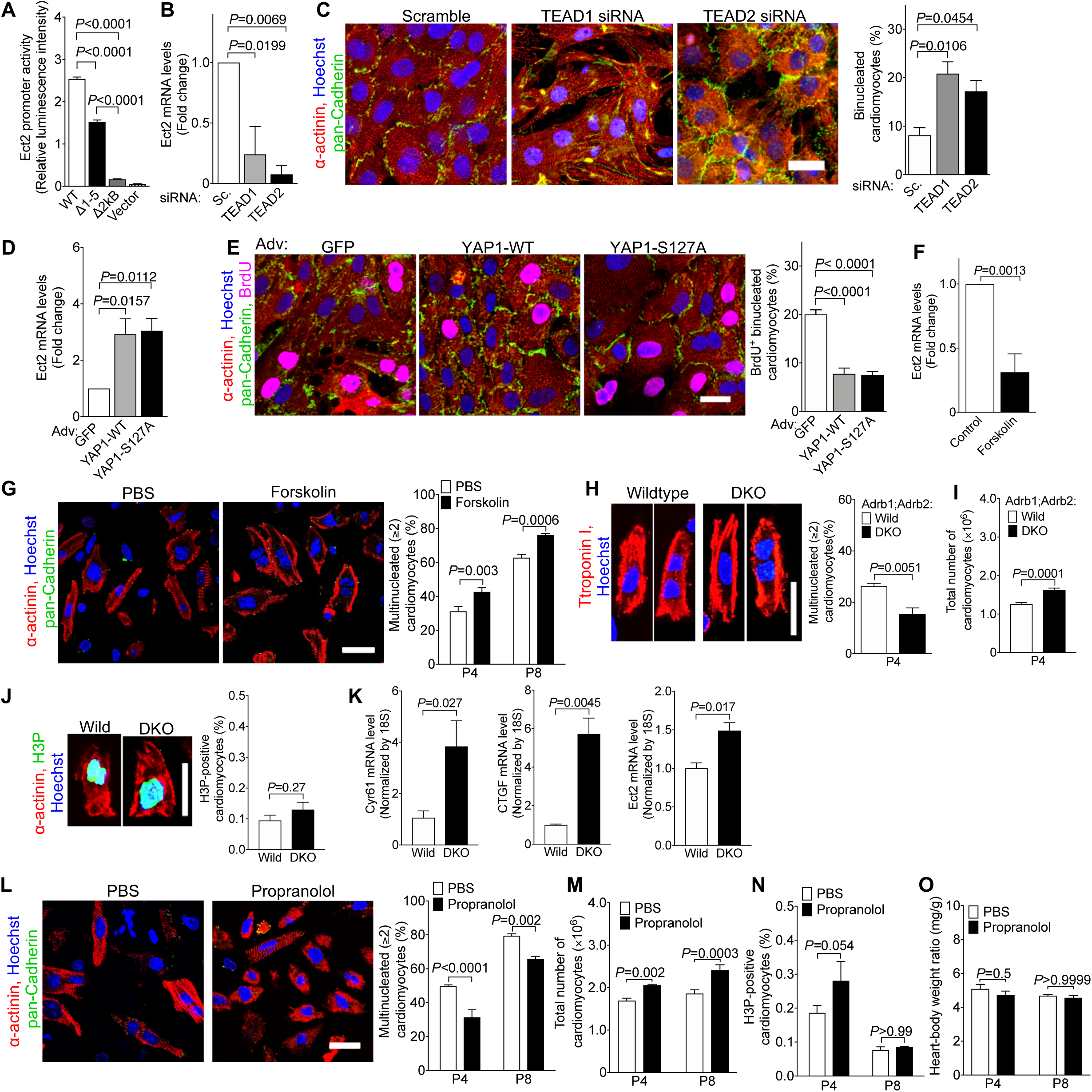
β-adrenergic receptor signaling regulates cardiomyocyte abscission and endowment in mice. (**A**) Removal of the five TEAD1/2-binding sites (**Fig. S7**) reduced the activity of the Ect2 promoter in luciferase assays in HEK293 cells. WT: wild type Ect2 promoter; Δ1-5: All five putative TEAD-binding sites were removed; Δ2kB: the continuous 2kB DNA sequence containing all five TEAD-binding sites was removed; Vector: Empty vector that did not contain Ect2 promoter (n = 4 cultures). (**B, C**) Knockdown of TEAD1 and TEAD2 by siRNA reduced Ect2 mRNA (**B**), and increased the proportion of binucleated NRVMs (P2, **C**, n = 3 cardiomyocyte isolations). (**D-E**) Adenoviral overexpression of wild type YAP1 (YAP1-WT) and a mutated version containing a S127A mutation (YAP1-S127A) in NRVMs (P2) increased the level of Ect2 mRNA (**D**) and reduced the percentage of binucleated cardiomyocytes (**E**, n = 4 cardiomyocyte isolations). (**F**) Forskolin reduced the mRNA level of Ect2 in cultured NRVMs (n = 5 cardiomyocyte isolations). (**G**) Forskolin administration (1 μg/g, 1 i.p. injection per day) increased the proportion of binucleated cardiomyocytes *in vivo* (n = 6 hearts/group). (**H**-**K**) Inactivation of β1- and β2-adrenergic receptor genes (DKO) decreased formation of multinucleated cardiomyocytes *in vivo* (**H**, n = 4 hearts/group), increased the total number of cardiomyocytes (**I**, n = 7 hearts for wild, n = 5 hearts for DKO), did not change M-phase (**J**, n = 4 hearts/group), and increased the expression of the YAP target genes Cyr61 and CTGF (**K**, n = 3 hearts/group). (**L-O**) Propranolol administration (10 μg/g, 2 i.p. injections per day) reduced the proportion of multinucleated cardiomyocytes (**L,** n = 6 hearts/group), and increased the number of cardiomyocytes (**M,** n = 6 heart/group for P4, n = 4 hearts/group for P8), but did not alter M-phase activity (**N**, n = 4 hearts/group) or heart-body-weight ratio (**O**, P4: n = 7 hearts for PBS, n = 6 hearts for Prop; P8: n = 4 hearts/group). Scale bar: 20 µm (**C, E, G, H, J, L**). Statistical significance was tested with one-way ANOVA with Bonferroni’s multiple comparisons (**A-E**), Student’s *t*-test (**F, H-K**), and two-way ANOVA with Bonferroni’s multiple comparisons test (**G, L-O**).

The Hippo pathway is activated by G protein coupled receptors (GPCR) *via* the stimulatory G protein, Gs ^28^. Accordingly, we treated cultured NRVMs with forskolin, a mimic of active Gs ^51^, which decreased Ect2 mRNA levels (**Fig. 3F**). We administered forskolin in newborn mice and found a 37% increase in the proportion of binucleated cardiomyocytes after 4 days and a 21% increase after 8 days (**Fig. 3G**). Because β1- and β2-adrenergic receptors (β1-, β2-AR) are the major Gs-activating GPCR in cardiomyocytes, we examined β1-AR^-/-^; β2-AR^-/-^ (double-knockout, DKO, ^52, 53^ pups. These mice showed a lower proportion of binucleated cardiomyocytes (**Fig. 3H**) and a higher endowment at P4 (**Fig. 3I**). Their cardiomyocyte M-phase activity was not changed (**Fig. 3J**). β1-AR^-/-^; β2-AR^-/-^ DKO hearts showed increased transcription of the Hippo target genes *Cyr61* and *CTGF*, as well as *Ect2* (**Fig. 3K**). We then administered propranolol, a blocker of β1- and β2-AR, in newborn mice. Propranolol decreased the proportion of binucleated cardiomyocytes by 21% after treatment from P1 to P4, and by 17% after treatment from P1 to P8 (**Fig. 3L**). This was associated with a 22% and 30% increase of cardiomyocyte endowment at P4 and P8, respectively (**Fig. 3M**), without a change of cardiomyocyte M-phase (**Fig. 3N**) or heart weight (**Fig. 3O**). These results show that reducing β-adrenergic receptor signaling enables abscission, thus increasing the endowment.

### Increasing cardiomyocyte endowment by administration of propranolol in neonatal mice improved heart function and reduced adverse remodeling due to myocardial infarction in adulthood

Although the propranolol-increased cardiomyocyte endowment did not alter cardiac function (**Fig. 4A**), a larger endowment should confer a benefit after large-scale cardiomyocyte loss, for example, after myocardial infarction (MI). We tested this by administering propranolol in the first week after birth and then inducing myocardial infarction in adult mice (**Fig. 4B**). We determined cardiac structure and function with MRI and histology (**Fig. 4B**). Twelve days after MI, mice with propranolol-induced endowment growth had an ejection fraction of 42%, compared with 18% in control mice (**Fig. 4C**). The thinned region of the LV myocardium after myocardial infarction was significantly smaller (**Fig. 4D**), and the relative systolic thickening was higher (**Fig. 4E**), indicating less adverse remodeling. Importantly, the region of myocardium affected by ischemia, visualized by late Gadolinium enhancement (**Fig. 4F**), and the scar size, determined by histology (**Fig. 4G**), were not different. Propranolol-treated hearts had a 30% higher cardiomyocyte endowment after MI (determined by stereology, **Fig. 4H**), in keeping with the increased endowment before MI. The heart weight was not changed (**Fig. 4I**), indicating that the higher endowment reduced the maladaptive hypertrophy, which drives adverse remodeling after MI. Taken together, these results demonstrate that rescuing cardiomyocyte cytokinesis failure with propranolol at the end of development reduces adverse ventricular remodeling in adult mice.

**Figure 4.**
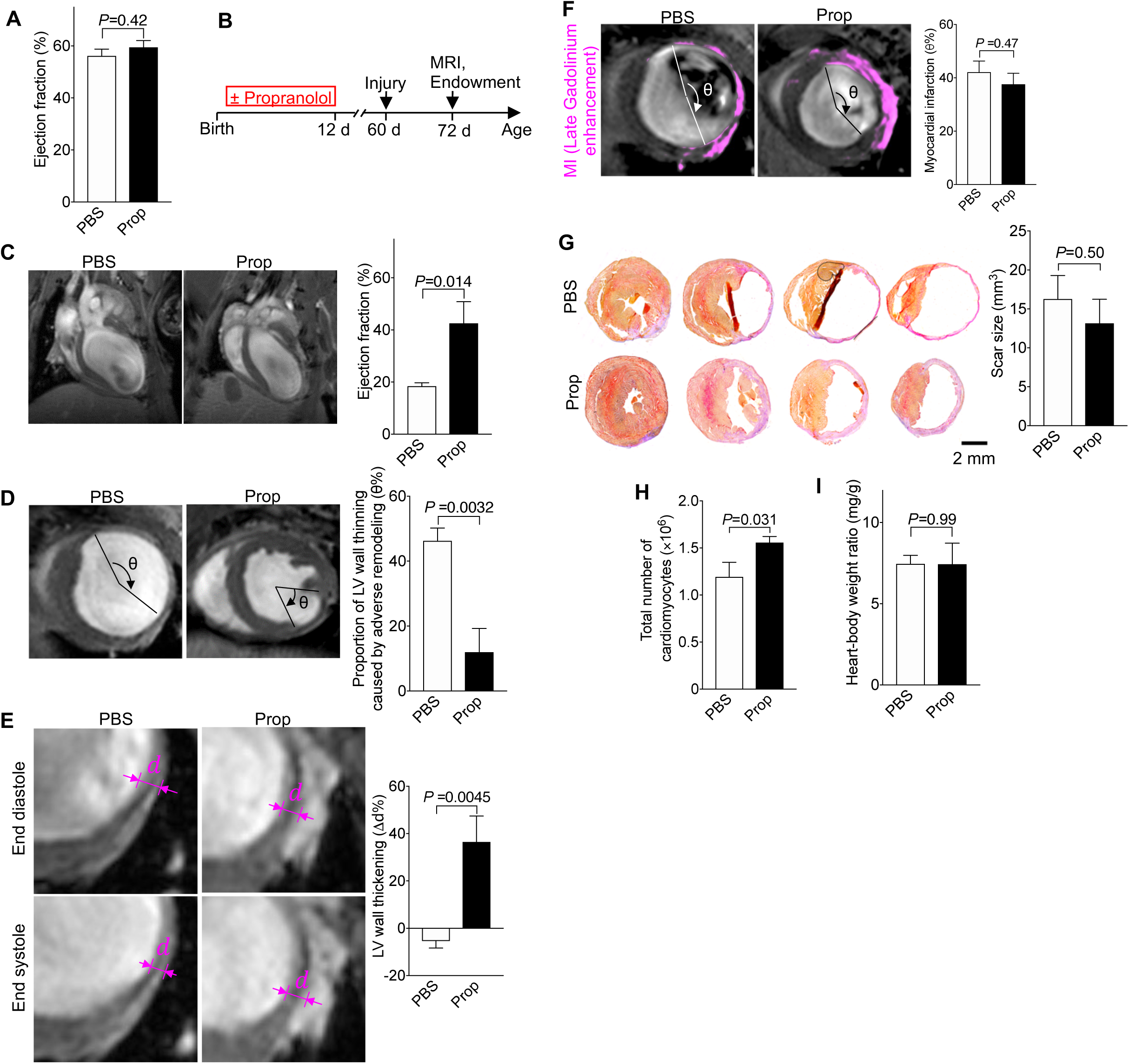
Mice with a 30%-increased cardiomyocyte endowment have better cardiac function and reduced adverse remodeling after myocardial infarction (MI). Mice received propranolol (Prop, 10 μg/g, 2 i.p. injections per day, P1-12). (**A**) The ejection fraction (EF) at P60 was not changed (n = 11 hearts for PBS, n = 6 hearts for Prop). (**B**) Diagram of experimental design. MI was induced at P60 and MRI and histology were performed 12 days after MI. (**C**) After MI, EF was higher in propranolol-primed mice. (**D-G**) The region of stretched-out myocardial wall is significantly smaller (**D**) and systolic myocardial thickening is greater (**E**), despite same scar size measured in vivo (**F**) and ex vivo (**G**, n = 5 hearts for PBS, n = 4 hearts for Prop). (**H, I**) Propranolol-primed hearts show a higher number of cardiomyocytes, determined by stereology (n = 5 hearts/group), after MI without change of heart weight (n = 5 hearts for PBS, n = 6 hearts for Prop) (**I**). Statistical significance was tested with *t*-test.

### β-blockers rescue cytokinesis failure in cardiomyocytes from infants with ToF/PS

We determined if the molecular mechanisms of cardiomyocyte cytokinesis failure we discovered are responsible for the increased proportion of binucleated cardiomyocytes in ToF/PS. To this end, we transcriptionally profiled single cardiomyocytes from infants with ToF/PS. ToF/PS cardiomyocytes showed lower Ect2 mRNA levels compared with dividing human fetal cardiomyocytes (**Fig. 5A**). The frequency of Ect2-expressing cycling cardiomyocytes in ToF/PS infants (24.4%) was significantly lower than in human fetal cardiomyocytes (75.6%, **Fig. 5B**).

**Figure 5.**
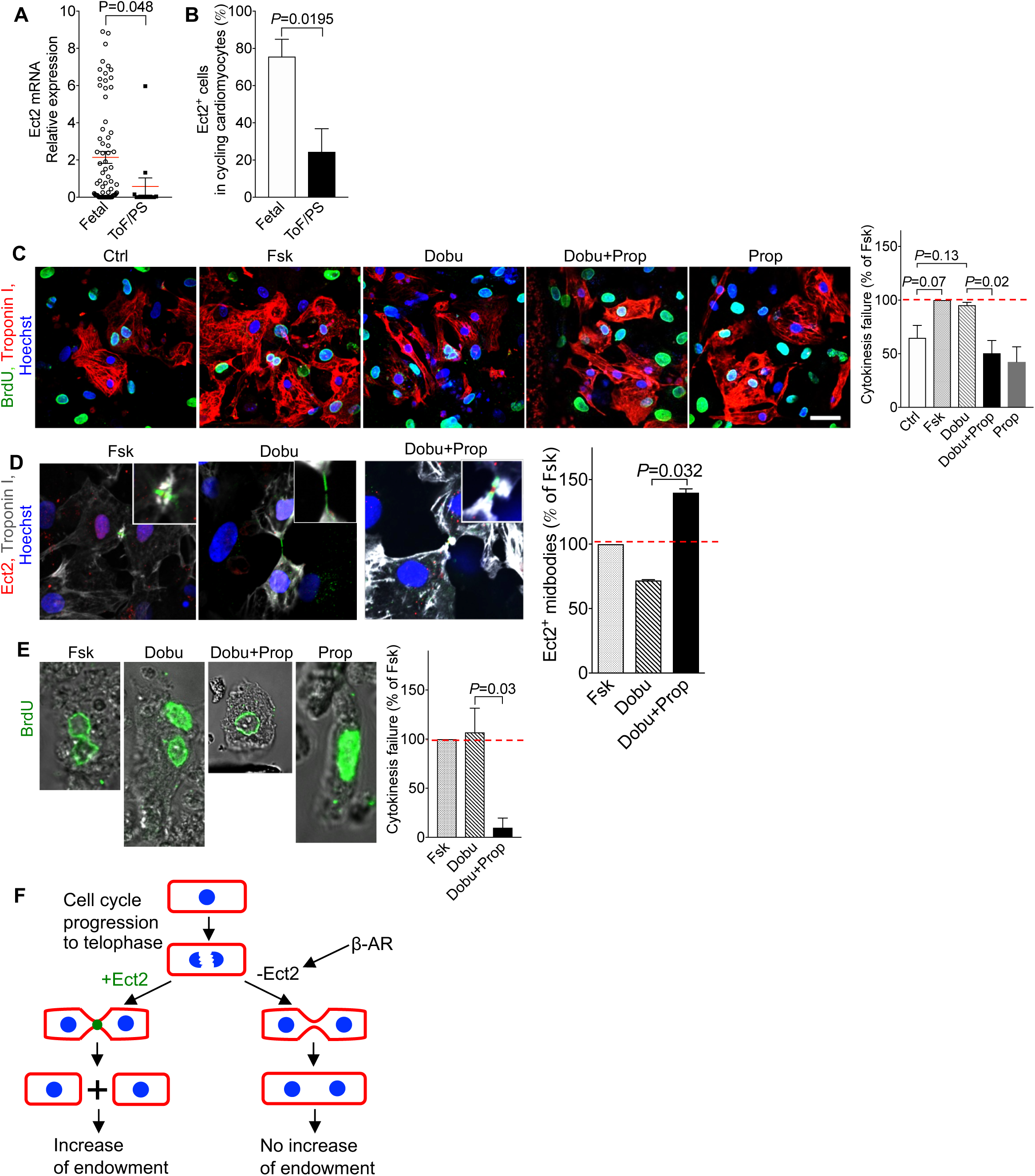
β-adrenergic signaling regulates cytokinesis in cardiomyocytes from ToF/PS patients. (**A**) Cardiomyocytes from ToF/PS patients exhibit decreased levels of Ect2 mRNA. Each symbol represents one cycling cardiomyocyte. (**B**) The percentage of Ect2-positive cycling cardiomyocytes was reduced in ToF/PS patients, compared with fetal human hearts. (**A**-**B**, Fetal: n = 71 cardiomyocytes from 4 hearts, ToF/PS: n = 13 cardiomyocytes from 3 hearts). (**C, D**) β-AR signaling regulates cytokinesis failure in cultured human fetal cardiomyocytes (n = isolations from 3 hearts; Ctrl: control; Fsk: Forskolin, Prop: Propranolol; Dobu: Dobutamine: 10 μM), measured by reduced formation of binucleated daughter cells (**C**) and higher prevalence of Ect2-positive midbodies (**D**). (**E**) Propranolol increases completion of cytokinesis in cultured ToF/PS cardiomyocytes (n = cultures from 3 patients). (**F**) Cellular model connecting cytokinesis failure to endowment changes. (**G**) Molecular model of cytokinesis failure in cardiomyocytes. Scale bar: 40 µm (**C**). Statistical significance was tested with T-test (**A**) and one-way ANOVA with Bonferroni’s multiple comparisons test (**C-E**).

Thus, cardiomyocytes in ToF/PS infants exhibit decreased Ect2 levels, similar to cardiomyocytes in neonatal mice (see **Fig. 2B**). This prompted us to examine the regulation by β-receptors. We used cultured human fetal cardiomyocytes and added forskolin to maximally increase cardiomyocyte cytokinesis failure (**Fig. 5C**). We then treated with dobutamine to mimic the *in vivo* microenvironment of increased β-AR stimulation, which increased binucleated cardiomyocytes to 95.2% of the forskolin-induced increase (**Fig. 5C**). Addition of propranolol blocked the dobutamine-stimulated increase of cardiomyocyte cytokinesis failure completely (**Fig. 5C**). We examined cardiomyocytes in cytokinesis by immunofluorescence microscopy, which showed that Ect2-positive midbodies were increased with propranolol (**Fig. 5D**). We then generated organotypic cultures of heart pieces from infants with ToF/PS and added BrdU to label cycling cardiomyocytes (**Fig. 5E**). Forskolin and dobutamine induced a maximal increase of binucleated cardiomyocytes, and propranolol inhibited the dobutamine-stimulated increase completely (**Fig. 5E**). In conclusion, β-receptors regulate cytokinesis failure in cardiomyocytes from infants with ToF/PS and propranolol decreases this effect.

## DISCUSSION

Our results show that ToF/PS infants develop increased cardiomyocyte terminal differentiation, measured by increased formation of binucleated cardiomyocytes. This happens in the first 6 months after birth, which shows that these changes do not result from surgical or medical interventions. The changes persisted in older ToF/PS patients. The increased binucleation indicates a proportionate failure of cytokinesis, reducing cardiomyocyte proliferation by 25%. By identifying the upstream molecular regulators, we show that β-blockers could turn cytokinesis failure to increased division.

The results in mice demonstrate that promoting the progression of cytokinesis to abscission in the post-natal period increases the endowment, which improves remodeling due to myocardial infarction. This raises the possibility that a decreased or increased cardiomyocyte endowment connects to outcomes in human patients. This could be tested with β-blocker administration in human infants with CHD to increase the endowment, followed by measuring clinical outcomes, such as myocardial function and risk of heart failure development. β-blockers have been used acutely to treat and prevent cyanotic spells in ToF/PS^54^. Although this demonstrates that β-blockers are safe in this population, an effect on myocardial growth mechanisms was not evaluated. Our results also predict that administration of β-blockers should produce the largest effect on cardiomyocyte cytokinesis in the first 6 months after birth, which is in line with our previously demonstrated effectiveness of stimulating cardiomyocyte cell cycling in CHD cardiomyocytes in the same period^55^. The duration of this period should also be assessed in types of CHD other than ToF/PS.

Elucidating the mechanisms generating binucleated cardiomyocytes allows us to compare this process in other cell types. Formation of binucleated cells is also an early event in cancer formation, leading, *via* entrapment of lagging chromosomes in the cleavage furrow, to aneuploidy ^56^. In this process, chromosome entrapment lowers Rho A activity in the cleavage furrow, leading to relaxation of the contractile ring and cytokinesis failure. However, we have no evidence for chromatin entrapment cardiomyocyte cytokinesis failure. In cancer cells, cytokinesis failure activates the Hippo tumor suppressor pathway^57^; however, we found that Hippo pathway activation triggers cytokinesis failure in cardiomyocytes. Cytokinesis failure is also a step in platelet formation, also induced by repression of Ect2 ^58^. This suggests a generalizable molecular mechanism of Ect2 repression in cytokinesis failure in somatic cells.

Cardiomyocyte cell cycle withdrawal in mice happens in the first 3 weeks^59^ and formation of binucleated cardiomyocytes in the first 2 weeks after birth^18, 19^. This is nearly coincident, which suggests that they could be mechanistically connected. However, we show that by increasing or decreasing cardiomyocyte binucleation, cardiomyocyte cell cycle entry does not change, thus demonstrating that formation of binucleated cardiomyocytes and cell cycle withdrawal are distinct molecular processes. This is supported by the increase of multinucleated cardiomyocytes (>2 nuclei) in ToF/PS, which shows that in ToF/PS, binucleated cardiomyocytes re-enter the cell cycle. This is consistent with the literature demonstrating that pig cardiomyocytes have up to 16 nuclei^60^, which shows that binucleated cardiomyocytes can re-enter the cell cycle multiple times.

Our results place the Hippo pathway downstream of β-AR signaling and upstream of *Ect2* gene regulation. This does not involve regulation of cell cycle entry and, as such, is distinct from direct genetic modulation of the central Hippo kinases and scaffolds, or regulation of the dystroglycan/agrin complex^27^, which all have a significant effect on cell cycle entry. Because of the large number of different G protein coupled receptors (GPCR), it is possible that other GPCR may have a function in regulating cardiomyocyte cell cycle entry, progression, and division.

Low levels of cardiomyocyte proliferation in mammals continue to be a barrier for heart regeneration ^27^. To overcome this, molecular interventions to stimulate cardiomyocyte cell cycle entry in adults have been proposed: increase of individual positive cell cycle regulators (cyclins A2 and D2, ^61, 62^ and combinations ^63^), removal of negative cell cycle regulators (p53, pocket proteins, ^64, 65^), and administration of mitogenic growth factors (FGF, NRG1, Oncostatin M, FSTL1, ^55, 66–68^). All of these approaches involve oncogenes or tumor suppressors and are associated with the risk of inducing uncontrolled proliferation. The findings presented here indicate that targeting the final stage of the cell cycle, *i.e.*, abscission, is a viable strategy that could synergize with any of these interventions to increase cardiomyocyte generation.

## Acknowledgments

We thank Tae Kyung Kim (University of Pennsylvania) for help with initial single cell amplifications. We thank Channing Der (University of North Carolina Chapel Hill) for providing Ect2^flox^ mice and Mark Petroncski (Clare Hall Laboratories, London), Buzz Baum (University College London), and Toru Miki (Nagaoka University of Technology) for providing Ect2 expression constructs. We thank Maria Magaro (Harvard Medical School) for technical assistance with phenotyping of cardiomyocytes. We thank the patients and families for participating in this research and the operating room staff and cardiac surgeons for assistance with identifying study subjects and ascertaining samples. We thank Sean Lal and Cris dos Remedios (University of Sydney) and Charles McThiernan (University of Pittsburgh) for providing human myocardial samples. This project used the UPMC Hillman Cancer Center and Tissue and Research Pathology/Pitt Biospecimen Core shared resource which is supported in part by award P30CA047904. We thank members of the Kuhn laboratory for support, helpful discussions, and critical reading of the manuscript.

## Author contributions

H.L., C.-H.Z., S.S., G.M.U., M.S., and B.K. developed the research strategy. S.S., G.M.U., M.A., and S.W. developed assays.

H.L., C.-H.Z., S.S., G.M.U., M.A., N.A., B.G., L.H., K.R., N.C., C.L., S.W., Y.W., D.Y., and S.C. performed experiments.

S.S., J.S., and S.C. performed transcriptional analysis of single mouse cardiomyocytes, with direct input and supervision by J.H.E.

N.A. performed mouse surgery, stereology, and transcriptional analysis of single human cardiomyocytes.

M.K., J.K., A.B., D.K., J.D., Z.B.-J. performed bioinformatics analysis of single-cell transcriptomes.

K.L., K.F., N.A., and B.K. developed the approach for identification of human study subjects and ascertaining of myocardium.

K.L. and K.F. identified human subjects.

M.S. and M.V. assisted K.L. and K.F. in identification of human study subjects and assisted N.A. in ascertainment of human heart samples.

M.L.S. and B.K. designed the human labeling and MIMS approach; C.G. and M.L.S. performed MIMS analysis.

H.L., C.-H.Z., S.S., G.M.U., M.A., N.A., C.L., and B.K. wrote parts of the manuscript, which

B.K. assembled, and all authors edited.

B.K. supervised and coordinated the entire study.

## Data availability

The datasets generated or analyzed for this study are available from the corresponding author upon request. The single-cell transcriptional profiling data have been deposited in NCBI’s Gene Expression Omnibus (GEO) with the dataset identifiers GSE108359 and GSE56638.

## Supplementary Materials

### MATERIALS AND METHODS

#### Study design and statistics

The study design, including the number of animals and the numbers of the cells counted, was predefined by the investigators. The genotypes of all pups were recorded throughout the entire period of the project. For the studies involving human tissue, the number of tissue samples was determined according to the availability of the samples. The investigators were blinded for the quantification of samples. The number of biological and technical replicates is provided in **Table S4.**

#### Approval of research involving animals and humans

Animal experiments were approved by the Institutional Animal Care and Use Committee (IACUC). Research involving human hearts was approved by the Institutional Review Board (IRB). Myocardium was resected as part of standard care.

#### Isolation of cardiomyocytes for culture

The Neomyts Cardiomyocyte isolation kit (Cellutron) was used to isolate cardiomyocytes from the fresh heart tissue (rat, mouse, and human) for cell culture study, according to the protocol provided by the vendor. The isolated rat and mouse cardiomyocytes were cultured on glass surface coated with fibronectin (10 μg/ml). The isolated human fetal/infant cardiomyocytes were cultured on glass surface coated with laminin (20 μg/ml).

#### Isolation of intact cardiomyocytes for quantification

We used a previously validated Fixation-Digestion method to isolate intact cardiomyocytes from fresh or frozen myocardium for quantification^1^. The fresh or frozen myocardium was cut into 1 mm sized tissue blocks followed by incubation with 3.7% formaldehyde at room temperature for 1.8 hours. After being washed with PBS 3 times to remove the formaldehyde, the fixed tissue blocks were digested in enzyme solution (Collagenase B, 3.6 mg/ml; Collagenase D, 4.8 mg/ml) at 37°C, with 10 rpm rotation. The digested cells were collected every 24 hours from the supernatant, and the undigested tissue was re-suspended with fresh enzyme solution until all of the tissue pieces were digested. All of the digested cells from the same heart were then transferred to the same tube and allowed to settle to the bottom of the tube. The cell pellet was collected for further study.

#### Mouse strains

##### Mouse strains expressing fluorescent cardiomyocyte marker and cell cycle reporter

Transgenic mAG-hGem mice were generated by Sakaue-Sawano et al ^2^. mAG-hGem consists of a green fluorescent protein, mAG 339 (monomeric Azami Green), fused to ubiquitination domains (110 amino acid residues at the N-340 terminus) of human DNA replication factor, serving as a cell cycle reporter in individual cells. As a result, cells with S/G2/M phase nuclei appear green. βMHC-YFP knock-in mice were provided by Oliver Smithies ^3^. We bred mice expressing both the mAG-hGem and βMHC-YFP constructs by crossing mAG-hGem homozygous mice with βMHC-YFP homozygous mice.

##### Inactivation of Ect2^flox^ gene in vivo, genotyping, and phenotyping

The *Ect2 ^flox^* gene was knockout using αMHC-Cre ^4^ and bred αMHC-Cre^+/-^;*Ect2^flox/+^* mice with *Ect2^flox/flox^* mice. To knockout the *Ect2 ^flox^* gene in newborn mice pups, we bred αMHC-MerCreMer^+/-^;*Ect2^flox/+^* mice. The newborn pups were administrated tamoxifen (30 µg/g body) through i.p. injection once daily from P0 to P2. The inactivation of *Ect2^flox^* gene was confirmed by PCR analysis of genomic DNA, as previously described. ^5^ The hearts were isolated, weighed, and imaged with a Leica dissection microscope equipped with a Nikon DSFi2 5 MP camera, or isolated for culture as described above.

##### Administration of propranolol and forskolin to ICR CD-1 mouse

Propranolol (10 µg/g body weight, 2 injections/day) and forskolin (1 µg/g body weight, 1 injection/day) was administered, respectively, to newborn mice from P1 to P8 via intraperitoneal (i.p.) injection. PBS was injected as the control group.

##### Timed pregnancies

Breeding was set up in the evening. Plugs were checked early the next morning and, if present, this was set as embryonic day 0.5.

#### Video microscopy

*To study the cleavage furrow regression in neonatal rat ventricular cardiomyocytes (NRVM),* the cardiomyocytes were isolated from neonatal rats (P2) were cultured in NS Medium for 48 hours prior to live cell imaging. The isolated cardiomyocytes were cultured in NS Medium containing 50 ng/μL NRG-1 (R&D Systems) in fibronectin-coated 35 mm glass bottom dishes (Part No. P35G-2-14-C-Grid, MatTek Corporation) or 8-well chambered coverglass (Part No. 155409, Lab-TekII). Cardiomyocytes were maintained in an environmental chamber (Tokai-HIT) fitted on the motorized stage (Prior) of an inverted Olympus IX81-ZDC autofocus drift-compensating microscope. Images were acquired at multiple positions every 30 minutes by a CCD camera (Hamamatsu) using an Olympus APON60XOTIRF objective, NA 1.49, together with differential interference contrast (DIC) components. Image acquisition and analysis were done using Slidebook™ 5.0 software.

*To characterize the effect of Ect2 overexpression*, neonatal rat cardiomyocytes (P2) were cultured in NS medium containing 10% FBS (Cellutron Life Technologies) for 3 hours to allow the cells to attach to the surface of an 8-well chambered cover glass coated with fibronectin (20 µg/ml). To remove unattached cells, the NS medium was changed carefully to DMEM/F12 (no phenol red) containing 5% FBS and adenovirus (Adv-Ect2-GFP, MOI = 2,000; Adv-GFP, MOI = 500). The chambered cover glass was mounted to an onstage incubator (Tokai Hit) providing a physiological environment (37°C, humidity, air mixture containing 5% CO2) on the motorized stage of a Nikon TiE microscope. Time-lapse imaging was performed for 72 hours. Images were acquired at 10-minute intervals by a CMOS camera (Andor Zyla) using a Nikon Plan Apo 60X oil objective, utilizing the Nikon perfect focus system (PFS) together with filter sets for observing GFP fluorescence. Image acquisition and analysis were done using Nikon NIS Elements 4.5 software.

#### Single mouse cardiomyocyte transcriptional profiling and data analysis

The freshly isolated cardiomyocytes from fetal (E14.5) and neonatal (P5) transgenic mAG-hGem mice were sorted on a FACSAria (20 psi, 100 μm nozzle, Becton Dickenson Biosciences). The cycling cells (mAG+) were separated from non-cycling cells (mAG-) based on the fluorescent signals (**Fig. S2**). The cardiomyocytes were identified by examining the expression of cardiomyocyte specific gene, *Tnnt2*, and non-cardiomyocyte contamination was identified and excluded by examining the expression of *Pdgfrb* gene (**Fig. S3**). Cardiac cells expressing mAG-hGeminin transgene (mAG+) were identified using a sequential gating strategy (**Fig. S2B-F**).

Initial size gates for forward scatter (FSC) *vs.* side scatter (SSC) were set to select the large cardiomyocytes corresponding to larger and more granular cells. Cell doublet discrimination was performed by a combination of high forward scatter height and area FSC-H/FSC-A and SSC-H vs. SSC-W plots. Live cells were selected by 7-aminoactinomycin D (7AAD, 1 μg/mL final concentration, Invitrogen) live/dead cell distinction staining. Finally, live 7AAD negative cells were distinguished by their mAG fluorescence intensity using the FACSAria 488-nm excitation laser. The mAG+ and mAG-cell fractions were collected separately for further downstream analyses. FACSDiva Software was used for data acquisition and analysis. The cycling cardiomyocytes (mAG-positive) were sorted into 96-well plates containing reverse transcriptase buffer for the following linear amplification. The freshly isolated single cardiomyocytes from the human myocardium collected during surgery (ToF/PS) and abortion (Fetal) were FAC sorted into 96-well plates containing reverse transcriptase buffer for the following transcriptional profiling and RNA sequencing. Following the synthesis of the first strand of cDNA, the molecular identity of the collected cardiomyocytes was confirmed by PCR for positive expression of the cardiomyocyte-specific gene Tnnt2 and negative expression of the non-cardiomyocytes gene Pdgfrb (**Fig. S3**). After 2 rounds of linear amplification (*in vitro* transcription), the RNA samples were sequenced using HiSeq 2500 (Illumina). To analyze the data of RNA sequencing, we first trimmed reads for adapters and poly(A) contamination using trimmomatic^6^. We then mapped the trimmed pair-end reads to human/mouse genome build hg19 using HISAT2 (ref. ^7^).

Using human/mouse gene annotations (.gtf) downloaded from iGenomes, we assigned read counts to genes using HTSeq and htseq-count^8^. Only exonic reads were counted with overlap assigned using the intersection non-empty method. We then further filtered all human samples by removing those with less than 60% overall reads mapping ratio and less than 1000 expressing genes. The remaining samples were used for further analysis. Finally, to mitigate the difference between samples (e.g. reads counts difference due to variable sequencing depth), we normalized the samples using the housekeeping genes. We started by normalizing the gene expression (exonic reads count) to total mapped reads in the sample. Next, we obtained a list of housekeeping genes with most stable expression in heart from the study as reference genes^9^. For each sample, we calculated the geometric mean of those reference genes. Then, we calculated the average of the geometric mean across all samples. This average was further divided by the geometric mean of the reference genes in each sample to get a sample-specific normalization factor. Multiplying the gene expression counts by the lane-specific normalization factor, we get the normalized expression. Lastly, we converted the normalized expression into log2 space.

#### Reverse transcription and real-time RT-PCR detection of mRNA levels

The mRNA was extracted from cardiac tissue samples using TRIzol reagent (Life Technologies) following the protocol provided by the vendor. Briefly, the samples were first homogenized using a mortar and pestle. The TRIzol reagent (1 mL) was added for each homogenized sample. Homogenized samples in TRIzol reagent were incubated at room temperature for 5 minutes, after which 0.2 mL of chloroform was added to each 1 mL of TRIzol. The mixtures were shaken by hand for 15 seconds and then incubated for 2-3 minutes at room temperature. The mixtures were then centrifuged at 12,000 x g for 15 minutes at 4°C. The clear upper aqueous phase containing the isolated RNA was carefully removed. The RNA was purified using the Qiagen RNeasy Plus Mini kit per the manufacturer’s instructions. The purified RNA was then quantified on a Nanodrop. Reverse transcription was performed on the purified RNA using the BioRad iScript cDNA synthesis kit (Catalog # 1708890). The kit was used according to the vendor protocol.

#### TaqMan assay for gene expression analysis

qRT-PCR was performed using Fast Taqman reagents (Thermo-Fisher). Probes were obtained from Thermo-Scientific including CYR61 (Mm00487498_m1), CTGF (Mm01192933_g1), ECT2 (Mm00432964_m1), and 18S (4319413E). All reactions were performed using a 1:10 diluted cDNA while mRNA expression levels were estimated using the 2DDCt method.

#### Ect2 mRNA expression in cultured NRVMs after forskolin treatment

The fresh isolated P2 NRVMs were treated with 10 µM forskolin for 3 days in culture. The Ect2 mRNA was quantified by qPCR using the method described above. The primers are listed in **Table S5**.

#### Adenoviral transduction, quantification of bi/multi-nucleation, and cell cycle activity

To overexpress Ect2 in primary cardiomyocytes, GFP-tagged full-length Ect2^10^ was cloned into pAd/CMV/V5-DEST^TM^ gateway vector (Thermo Fisher, Cat#: V49320) and adenoviruses were generated in 293A cells according to the manual’s instructions. The isolated cells were cultured for 12 hours, washed gently with culture medium, and then cultured in the culture medium containing adenovirus (Adv-Ect2-GFP, MOI = 2000; Adv-GFP, MOI = 500) and BrdU (30 µM). After being cultured for 48 hours, the medium was replaced by fresh culture medium containing BrdU (30 µM) without adenovirus (48 hours). The cells were then fixed at day 5 and stained for BrdU and pan-cadherin or H3P. The binucleation of BrdU-positive cardiomyocytes and H3P-positive cardiomyocytes was quantified.

#### Examination of TEAD-binding sites by Luciferase Assay

##### Construct preparation

To generate deletions of the Ect2-promoter region, we first subcloned the 2.8 kb Ect2 promoter region into the luciferase reporter construct pGL4.10 (Promega, Madison WI). Using this construct, we generated a deletion construct that removed all five TEAD-binding sites and the DNA between them (Ect2-Promoter – Δ2kb) and a construct that only removed the nucleotides of the binding sites (Ect2-Promoter-Δ1-5). All plasmid constructs were amplified using TOP-10 competent bacteria and plasmid DNA was isolated and purified using the Qiagen Midi-Prep System (Qiagen, Germany) according to the manufacturer’s instructions. Once purified, the plasmid constructs were sequence-verified to eliminate the possibility of unwanted mutations. *Transfection into HEK-293 cells:* To measure the ability of the Ect2-promoter regions to generate a luciferase signal, we transfected HEK-293 cells with the following plasmids: 1) wt-Ect2-promoter-pGL4.10; 2) Ect2-Promoter – Δ2kb – pGL4.10; 3) Ect2-Promoter-Δ1-5-pGL4.10; 4) Empty – pGL4.10; 5) pGL3.1-Renilla-Control. 4 µg of plasmid DNA were transfected into one well of a 6-well plate of HEK-293 cells using the lipofectamine 2,000 system (Invitrogen, CA) according to the manufacturer’s instructions. Each pGL4.10 group was transfected along with an equal amount of the pGL3.10-Renilla-Control vector.

Following transfection, the cells were allowed to incubate in a standard tissue culture incubator for 48 hours to allow for optimal luciferase construct expression. *Quantification of Ect2 promoter activity:* To perform the luciferase measurements on cells transfected with Ect2-promoter deletions, we used the Dual Luciferase Reporter Assay System #E1910 (Promega, Madison, WI). After 48 hours of incubation, the media was removed and the cells were washed 2 times with PBS. Then the cells were lysed through incubation in the supplied passive lysis buffer for 20 min. After lysis, the mixture was centrifuged at 13,000 x g for 5 minutes to pellet the debris, and the cell lysate was collected for subsequent analysis. The luciferase activity in each group was quantified according to the manufacturer’s instructions. We prepared solutions of the Stop-n-Glo reagent and the Luciferase Assay II reagent (LAR2) as described in the manufacturer’s instructions. The prepared solution was then loaded into the injectors of a Synergy H1 Hybrid plate reader (Biotek, Winooski, VT), and the first 100 µl of the solution was dispensed. We then placed 20 µl of cell lysate into the bottom of a Greiner Cellstar™ microclear bottom 96-well plate (Sigma, St. Louis, MO) and loaded it into plate reader. 100 µl of LAR2 was then dispensed into 1 well and measurement of firefly luciferase was taken over a period of 10 sec. After measurement, 100 µl of Stop-n-Glo reagent was injected into the same well and incubated for 5 sec to quench the firefly luciferase reaction. The measurement of Renilla-control luciferase was performed for 10 seconds to measure the background luminescence activity. The measurement was repeated for each well. The difference between luminescence obtained from the experimental firefly luciferase and the Renilla luciferase measurements was calculated as the activity of the Ect2 promoter. The cell lysate from each transfection was transferred to 10 wells of a 96-well plate. The values of the activity of the transfected Ect2 promoter were averaged among the 10 wells.

#### Immunocytology and Immunohistology

*To study the cultured cardiomyocytes attached to the surface of substrate,* the cultured cardiomyocytes were fixed with 3.7% formaldehyde or 4% paraformaldehyde for 12 minutes at room temperature. After being washed in PBS 3 times, the cells were immersed in permeabilizing and blocking solution (0.5% Triton X-100, 5% donkey or goat serum in PBS) for 30 minutes. Then the mixture of primary antibodies was added to the samples and incubated overnight at 4°C. After being washed 3 times in PBS, the cells were immersed in a mixture of secondary antibodies and incubated for 1 hour at room temperature. The nuclei were counterstained with Hoechst 33342 (Invitrogen, dilution 1:1000) for 5 minutes at room temperature. Then, the samples were dipped into distilled water for 15 seconds. The cells were mounted in 10 µl mounting media containing 1% N-propyl-gallate dissolved in glycerol and sealed with nail polish.

*To study the activity of RhoA (RhoA-GTP) at the midbody*, the cells were fixed in 0.5 ml ice-cold trichloroacetic acid (TCA) solution (10% w/v) for 15 minutes and then washed 3 times with PBS containing 30 mM glycine^11, 12^. The cells were incubated in an ice-cold mixture of permeabilizing and blocking solution (5% goat serum, Triton X-100 0.5 µl/ml in PBS) for 60 minutes. After being washed with PBS containing 30 mM glycine 3 times, the cells were incubated in a mixture of primary antibodies (RhoA-GTP, mouse IgM, New East Biosci, 26904, dilution 1:100); α-actinin (mouse IgG1, Sigma A7811, dilution 1:200); Aurora B kinase (rabbit, Abcam, ab2254, dilution 1:200) was added to the samples (50 µl/coverslip) and incubated overnight at 4°C. After being washed with ice-cold PBS containing 30 mM glycine 3 times, a mixture of secondary antibodies (goat anti-mouse IgG1 633, ThermoFisher A-21126, dilution 1:200; goat anti-mouse IgM 594, ThermoFisher A-21044, dilution 1:200; goat anti-rabbit 488, ThermoFisher A-11034, dilution 1:200) was added to the samples (50 µl/coverslip) and incubated for 60 minutes at room temperature. The nuclei were counterstained with Hoechst 33342 (Invitrogen, dilution 1:1,000) for 5 minutes at room temperature. Then the samples were dipped into distilled water for 15 seconds. The cells were mounted in 10 µl mounting media containing 1% NPG dissolved in glycerol and sealed with nail polish.

*To stain the Fixation-Digested cardiomyocytes in suspension*, if BrdU staining was necessary, the isolated cardiomyocytes were first incubated in 2N HCl solution containing Triton X-100 (Fisher Scientific, BP 151-500) for 60 minutes at room temperature. The HCl was then neutralized with the same amount of 2N NaOH solution. The isolated cardiomyocytes were then blocked in a solution of 5% goat serum (Sigma, G9023-10 ML), or donkey serum (Sigma, D9963-10ML), and incubated in primary antibodies at 4°C overnight. After PBS wash, the cells were incubated in mixture of secondary antibodies at 4°C overnight. The cells were washed with PBS, and incubated in Hoechst solution (PBS, 1:1000) at room temperature for 5 minutes followed by three rounds of PBS wash. The cells were then ready for future study.

*To study the hearts using immunohistology*, the hearts were resected and fixed immediately in 3.7% formaldehyde at room temperature for 8 hours. The fixed hearts were washed in PBS and immersed in 30% sucrose solution at 4 °C for 24 hours. After being embedded in optimum cutting temperature (OCT) compound, the frozen block was trimmed and the exposed heart tissue was cut into 20 μm thick sections with a Leica CM1950 cryostat and adhered to glass slides (Color Frost, Fisher). The slides were fixed in 3.7% formaldehyde, permeabilized in 0.5% Triton X-100, and blocked in PBS containing 20% goat serum and 0.2% Tween 20. Then the mixture of primary antibodies was added to the samples and incubated overnight at 4°C. After being washed 3 times in PBS, the cells were immersed in a mixture of secondary antibodies, and incubated for 1 hour at room temperature. The nuclei were counterstained with Hoechst 33342 (Invitrogen, dilution 1:1000) for 5 minutes at room temperature. Then, the samples were dipped into distilled water for 15 seconds. The cells were mounted in 10 µl mounting media containing 1% N-propyl-gallate dissolved in glycerol and sealed with nail polish.

#### Quantification of mouse and rat cardiomyocyte cell cycle activity and proliferation

*To quantify the midbodies of the neonatal rat ventricle cardiomyocytes (NRVMs)*, we cultured the NRVM (E14.5 and P2) in NS medium for 3 days, then studied the midbodies using immunocytology. We first identified the cleavage furrows of the cycling cells at late cytokinesis stages, i.e. telophase, by visualizing the immunostaining for Aurora B kinase. We then identified cardiomyocytes using the markers troponin I and α-actinin, and examined the presence of the Ect2, Racgap 1, or RhoA-GTP at the midbodies.

*To count the total number of cardiomyocytes in a mouse heart,* all the digested cardiomyocytes from the same heart were suspended in 2 ml PBS buffer. The concentration of the cardiomyocytes inside the PBS buffer was measured using a hemocytometer. The total number of cardiomyocytes was then calculated by multiplying the measured cell concentration by the volume of the cell solution (2 ml).

*To quantify the binucleation and H3P+ cardiomyocytes in mouse hearts*, cardiomyocytes were isolated by fixation-digestion prior to immunostaining. The solution containing the stained cells was then transferred to 8-well chambered cover glass (400 µl/well). After the cells were settled down to the bottom of the chambers, the number of mono/bi/multinucleated cardiomyocytes were counted with a fluorescence microscope, and the percentage of mono/bi/multinucleated cardiomyocytes was calculated. To quantify the proportion of H3P-postive cardiomyocytes, the concentration of stained cardiomyocytes in suspension (*C_cardiomyocytes_*) was first measured using a hemocytometer. The cell suspension (400 μl) was then transferred to a well of a chamber slide, and the total number of H3P-positive cardiomyocytes (H*_H_*_3*P*+_) inside the well was counted with a TiE epifluorescence microscope. The proportion of the H3P-positive cardiomyocytes (P*_H_*_3*P*+_) was calculated using the following formula: 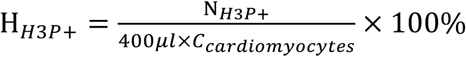.

*To measure the size (2D) of cardiomyocytes in mouse hearts,* the cells were imaged using a Nikon TiE microscope with a Zyla CMOS camera. The image layers from the immunofluorescence channels were used to identify cardiomyocytes and the layer from the bright-field channel was used to measure the area of single cardiomyocytes using ImageJ software.

*To quantify the bi- and multinucleated cardiomyocytes in mouse hearts,* the hearts from the αMHC-Cre;Ect2flox mouse (P1) and the ICR mouse (P4 and P8) were treated with propranolol, forskolin, and PBS were resected and the cardiomyocytes were dissociated using Fixation-Digestion method and immunostained. The mono/bi/multi-nucleated cardiomyocytes were then quantified under Nikon TiE microscope. About 500 cardiomyocytes in each sample were quantified.

#### Mouse left anterior descending (LAD) coronary artery ligation as a model of myocardial infarction

Newborn ICR mice were first treated with propranolol (10 µg/g, 2 i.p. injections/day) and PBS, respectively, from P1 to P12. To induce myocardial infarction at 8 to 12 weeks, we permanently ligated the left anterior descending (LAD) artery. Briefly, the mice were anesthetized with 4% isoflurane and then intubated and ventilated with a stroke volume of 225 µl at 145 breaths/minute. The hearts were exposed by performing thoracotomy and the LAD artery was tied with 8-0 nylon suture. After the chest was closed with 6-0 suture, bupivacaine (8µg/g) was locally administered through subcutaneous injection. The mice were kept intubated without isoflurane for another 30 minutes at room temperature and then at 37°C until waking up from the anesthesia. The bupivacaine (8µg/g) was kept administered through subcutaneous injection in the following 3 days (1 dpi to 3 dpi).

#### Stereology

The LAD ligated mouse hearts were isolated, gently washed in PBS, weighed, immersed in cardioplegia solution, and fixed overnight with 3.7% formaldehyde at room temperature. The hearts were then placed in 30% sucrose for 48 hours at 4°C. The atria for each heart were cut off and the ventricles were embedded in OCT.

*The hearts were sectioned on Leica CM1950 cryostat in cross-sectional orientation* (thickness 15 μm), resulting in 75-80 slides/heart with 4 sections each. For random systematic sampling we used a random number generator ranging between 1-6 to determine the first slide and then selected every 17^th^ slide for staining. The tissue sections were stained with *α*-actinin (sarcomeric stain) and Hoechst (nucleus stain).

The adjacent slides of *immunofluorescence-stained* slides were selected for *AFOG staining* and followed the standard lab protocol method. Photomicrographs were taken on Nikon TiE microscope (objective lens 10x).

*Quantification of myocardial volume by point count method on AFOG-stained sections:* Tissue sections were imaged on Leica MZ26 dissector microscope, zoom factor: 1.0X. The area of myocardium (red after AFOG staining) and scar (blue after AFOG staining) was quantified using ImageJ 1.51s version. To this end, the image was overlaid with a grid to determine area per point. The distance between selected sections was calculated (n^th^slide × #of sections/slide × section thickness). The LV volume was measured by counting the number of grids on both LV myocardium and the scar region. The scar size was also measured using the same method.

*Optical dissector method to determine the volume density of cardiomyocyte nuclei*: The immunofluorescence stained sections were imaged using Nikon A1R confocal microscope. We selected four random spots on the technically-best section from each slide and analyzed 20 random samples from each heart. The total number of positive cardiomyocyte nuclei per heart was counted and the mean per sample volume was calculated.

*Total number of cardiomyocytes (endowment):*

Number of cardiomyocytes = Number of cardiomyocyte nuclei / (Mono% + 2×Bi% = 3×Tri% + 4×Tetra%)

#### *In vivo* Cardiac MRI

##### Anesthesia

Mice were anesthetized with 4% isoflurane mixed with room air in an induction box for 1 to 3 minutes. The depth of anesthesia was monitored by tow reflex, extension of limbs, and spine positioning. Once the plane of anesthesia was established, the mouse was placed on a manually built neonatal mouse holder and the anesthesia was maintained by 1.5 to 2 % isoflurane with 100% oxygen via a nose cone. The respiration was continuously monitored by placing a small pneumatic pillow under the animal’s diaphragm which was connected to a magnet-comparable pressure transducer feeding to a physiological monitoring computer equipped with respiration-waveform measuring software (SA Instruments, Stony Brook, NY). The respiration waveform was automatically processed to detect inspiration, expiration and respiration rate.

##### In vivo CMR Acquisition

*In-vivo* cardiac MRI (CMR) was carried out on a Bruker Biospec 7T/30 system (Bruker Biospin MRI, Billerica, MA) with the 35-mm quadrature coil for both transmission and reception. Free-breathing-no-gating cine MRI with retrospective navigators was acquired with the Bruker Intragate module.

##### Late-gadolinium enhancement (LGE) for myocardial infarct evaluation

Subcutaneous injection of Multi Hance (Gadobenate dimeglumine, 529 mg/ml, Bracco Diagnostics, Inc, Monroe Twp, NJ 08831) was administered right before the CMR acquisition with the dosage of in 0.1 mmol Gd/kg bodyweight. T1-weighted images to highlight LGE was acquired 15-20 minutes after the subcutaneous administration of Multi Hance. Eight T1-weighted short-axis imaging planes covering the whole ventricular volume with no gaps were acquired with the following parameters: Field of view (FOV) = 2.5 cm × 2.5 cm, slice thickness = 1mm, in-plane resolution = 0.97 µm, flip angle (FA) = 10 degrees, echo time (TE) = 3.059 msec, repetition time (TR) = 5.653 msec.

##### Cine CMR to determine cardiac function

White-blood cine movies with 20 cardiac phases were acquired for each mouse with equivalent temporal resolution for the cine loops was about 16.5-21.5 ms per frame. Eight short-axis imaging planes covering the whole ventricular volume with no gaps and one long-axis plane were acquired with the following parameters: Field of view (FOV) = 2.5 cm × 2.5 cm, slice thickness = 1mm, in-plane resolution = 0.97 µm, flip angle (FA) = 30 degrees, echo time (TE) = 1.872 msec, repetition time (TR) = 38.293 msec.

#### Analysis of CMR results

##### Extent of myocardial infarction (MI) by late-gadolinium enhancement (LGE)

The extent of myocardial infarction was defined by the percentage of the myocardium displaying hyperintensity 15-20 minutes after Gd administration. To obtain the proportion of myocardial infarction (MN), we measured the angle of the portion of the myocardium displaying hyperintensity in the left ventricle wall of each scanned slice (O) and divided the angle by 360° to get the percentage of infarction of the slice (*MI_i_*). The infarction of each scanned slice was then averaged to have the proportion of myocardial infarction *MI* = ∑*_i_ MI_i_*.

*Cardiac function from cine CMR.* The left ventricular endocardium and epicardium boundaries of each imaging slice at the end-systole (ES) and the diastole (ED) were manually traced by a blinded operator on the Paravision 5.1 Xtip software (Bruker Biospin MRI, Billerica, MA) to calculate the following functional parameters: left ventricular blood volume (LVV), left ventricular wall volume (LV wall), LV mass, stroke volume (SV), ejection fraction (EF), heart rate (HR), cardiac output (CO), longitudinal shortening, and radial shortening. LVV is calculated by summation of all the short-axis slices. The ejection fraction (RS) was calculated

using the following equation: 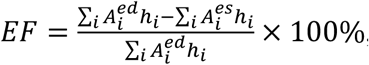 where 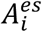 is the internal left ventricle area of slice O at end systole 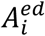, the internal left ventricle area of slice *i* at end diastole, and *h_i_* is the thickness of each scanned slice. The proportion of left ventricle wall thinning

caused by adverse remodeling was calculated by measuring the proportion of the angle of the thinned left ventricle caused by adverse remodeling after LAD ligation. To calculate the left ventricle wall thickening, we first selected the scanned slice that demonstrated the largest LGE portion. The thickness at both ends and the middle of the LGE portion was measured at end systole (*d^es^*) and diastole (*d^ed^*) and calculated the average. The left ventricle wall thickening was calculated through equation *LV wall thickening* = (*d^es^* _−_*d^ed^*)/*d^ed^* × 100%.

#### Examination of heart function of newborn mice by echocardiography

We performed echocardiography using a Vevo 770 device (VisualSonics) with a 25 MHz probe (RMV-710B). Two-dimensional (2D) B-mode recordings covering both ventricles were obtained in the left parasternal short axis view. The left ventricular endocardium and epicardium boundaries of the imaging slice at end-systole and diastole were manually traced by a blinded operator on the ImageJ software. The ejection fraction (RS) was calculated using the following equation: 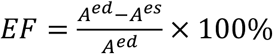 where *A^es^* is the left ventricular blood area at end systole, *A^ed^* is the left ventricular blood area at end diastole.

#### Quantification of cytokinesis in human cardiomyocytes

*To quantify the bi- and multinucleation of intact human cardiomyocytes,* the cardiomyocytes in the fresh or frozen right ventricle myocardium collected as part of standard care during surgical repair of the ToF/PS hearts were isolated through Fixation-digestion method, followed by immunostaining as described above. The cells were then imaged and quantified using Nikon A1R microscope for mono/bi/multi-nucleation. The cardiomyocytes isolated from the myocardium without heart disease were used as the control group.

*To quantify the response of human fetal cardiomyocytes to β-adrenergic receptor signaling reagents,* the human fetal cardiomyocytes were isolated from fresh specimen using the Cellutron Kit. The cells were cultured in IMDM media containing 10% FBS and recombinant neuregulin 1 (rNRG1, 100 ng/ml) for 2 days. The cells were then treated with forskolin (10 µM), propranolol (10 µM), dobutamine (10 µM), or a combination of propranolol (10 µM) and dobutamine (10µM) for 3 days, in the presence of BrdU (30 µM). The cells were then fixed with 3.7% formaldehyde and stained using primary antibodies against troponin I (goat, Abcam), pan-cadherin (rabbit, Abcam) and BrdU (rat, Abcam), and secondary antibodies from donkey against goat, rabbit, and rat IgG, respectively. The proportion of binucleated cardiomyocytes among the BrdU-positive cardiomyocytes was quantified with a Nikon A1R microscope.

*To quantify the response of ToF/PS myocardium to β-adrenergic receptor signaling reagents,* the myocardium samples were collected during ToF/PS correction and were cut into ∼1 mm tissue blocks and cultured in IMDM media containing 10% FBS, recombinant neuregulin 1 (rNRG1, 100 ng/ml), and BrdU (30 µM), for 6 days. During culture, the samples were treated with forskolin (10 µM), propranolol (10 µM), dobutamine (10 µM), or a combination of propranolol (10 µM) and dobutamine (10 µM). Cardiomyocytes were isolated with the Fixation-Digestion method. After denaturing the DNA with 2N HCl, and neutralization with 2N NaOH, immunofluorescence antibody labeling was performed in solution. The proportion of binucleated and BrdU-positive was quantified with a Nikon A1R microscope.

#### Quantification of cardiomyocyte binucleation in ToF/PS patients by Multi-isotope Imaging Mass Spectrometry (MIMS)

The clinical study protocol was approved by the IRB, and informed consent was obtained from the parents. Based on prior human MIMS studies ^13, 14^, the patient received ^15^N-thymidine (50 mg/kg p.o., Cambridge Isotope Laboratories, on five consecutive days) at 3.5 weeks of age. The patient underwent surgery at 6 months of age. A discarded piece of right ventricular myocardium was obtained, fixed in 4% paraformaldehyde, embedded in LR White, and 500 nm sections were mounted on silicon chips. Multiple isotope imaging mass spectrometry (MIMS) was performed on myocardial sections utilizing the NanoSIMS 50L (CAMECA) and previously described analytical methods ^15^ ^15^N-thymidine labeling was measured by quantification of the ^12^C^15^N^-^/^12^C^14^N^-^ ratio (**Fig. S8**), obtained in parallel with mass images utilized for histological identification (^12^C^14^N^-^, ^31^P^-^, ^32^S^-^). Quantitative mass images were then analyzed using OpenMIMS version 3.0, a customized plugin to ImageJ (NIH) that is available at https://github.com/BWHCNI/OpenMIMS ^16^. An observer blind to the ratio images identified nuclei and assigned cellular identity using the ^14^N (^12^C^14^N-), ^31^P-, and ^32^S-images as previously described ^15^. Cardiomyocyte nuclei were identified by their close association with sarcomeric structures.

#### Statistical analyses

Statistical testing was performed with Student’s t-test, Fisher’s exact test, and ANOVA, followed by Bonferroni post hoc testing, as indicated. A two-sided P value ≤ 0.05 was accepted as statistically significant. Statistical analyses were performed with GraphPad Prism, version 6.

---

**Figure S1.**
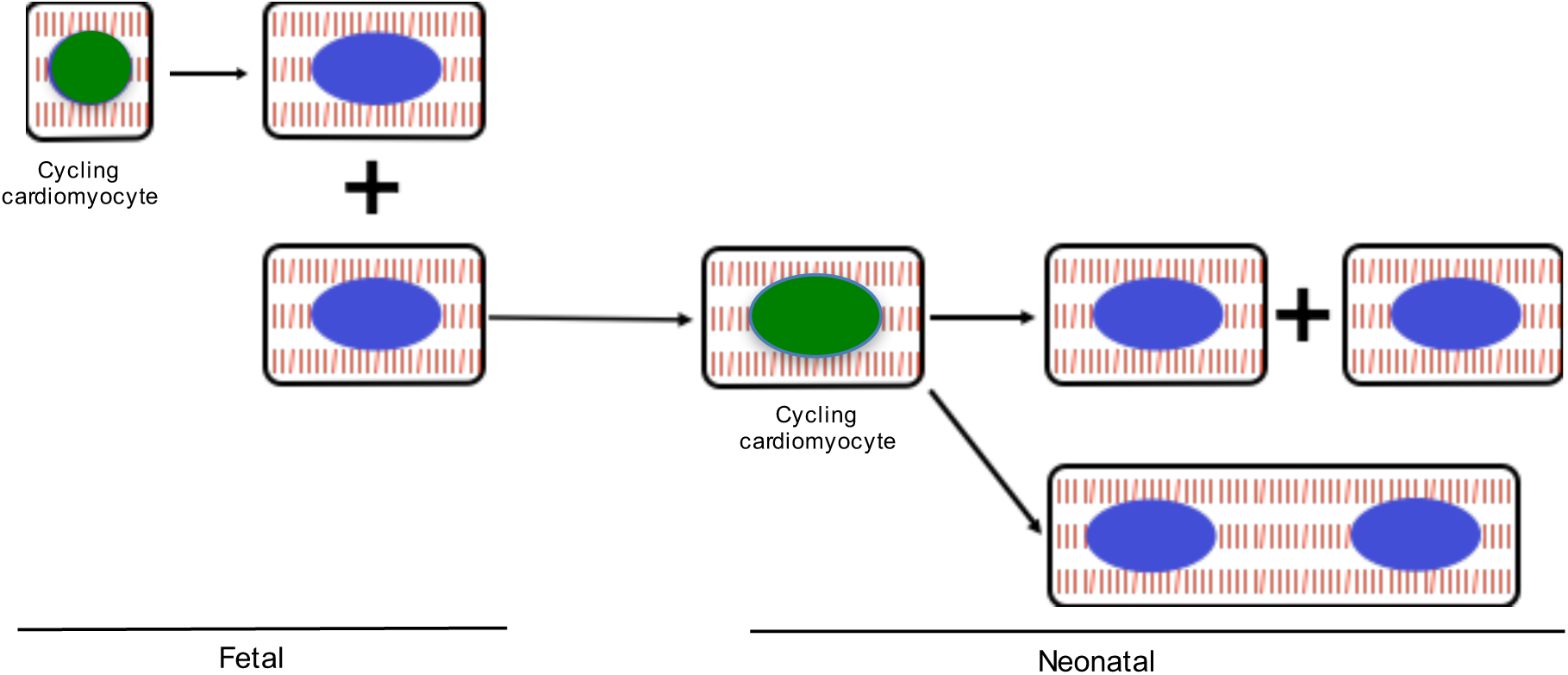
A single-cell transcriptional profiling strategy identifies molecular mechanisms of cardiomyocyte binucleation. When neonatal mouse cardiomyocytes are in the cell cycle, two outcomes are possible: the cardiomyocyte may either divide or become binucleated. As such, bulk RNAseq of cycling neonatal mouse cardiomyocytes would not reveal the molecular mechanisms that are specific for binucleation. To overcome this challenge, we have taken a single cell transcriptional profiling approach, which should isolate the specific molecular mechanism. We identified cycling cardiomyocytes with the Azami green-Geminin (mAG-hGem) reporter and isolated single cycling and not cycling cardiomyocytes with FACS, as demonstrated in **Figure S2**.

**Figure S2.**
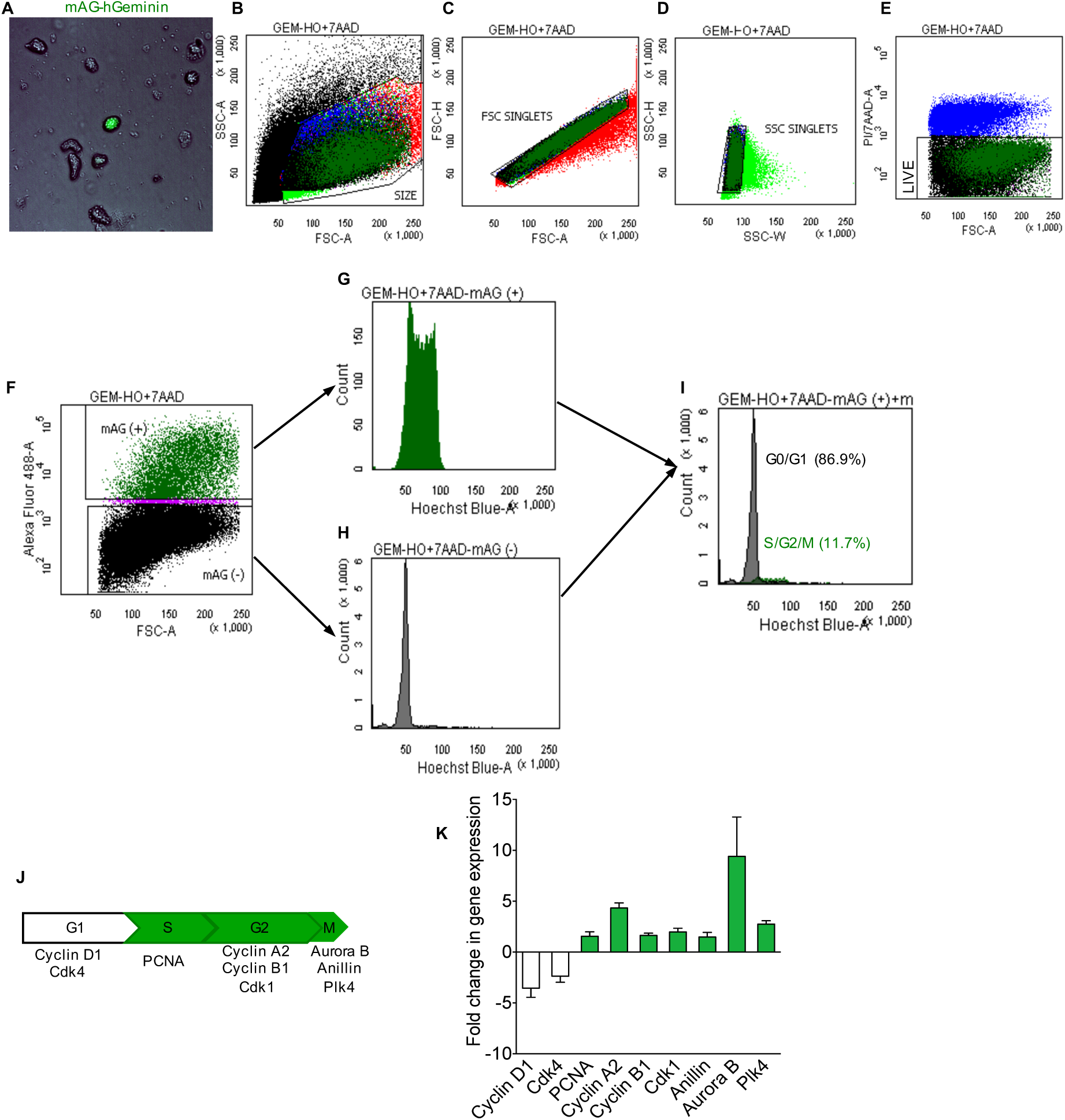
Strategy for isolation and characterization of cycling cardiac cells. (**A**) Cells were isolated from mAG-hGeminin E14.5 hearts and subjected to FACS. Panels (**B-F**) show FACS gating strategy for isolation of single cycling and non-cycling cardiac cells. We validated that the population of Geminin-positive cells were in the cycle by (**G-I**) analyzing DNA contents after staining with Hoechst, and (**J, K**) RT-PCR with primers for cell cycle genes. Cells were gated sequentially for size (**B**), doublet exclusion (**C, D**), viability (**E**), and mAG-hGeminin expression (**F**). Representative histograms of cycling Geminin-positive cells (**G**), non-cycling Geminin-negative cells (**H**), and merge of cycling Geminin-positive cells and non-cycling Geminin-negative cells show that Geminin-positive cells have higher DNA content (**I**). (**J**) Linear cell cycle diagram with marker genes selected for identification of cell cycle phases indicated below. The phases of the cell cycle indicated by mAG-hGeminin (S-, G2- and M-phase) are highlighted in green. (**K**) RT-PCR analysis of Geminin-positive cells shows no expression of the G1 phase markers cyclin D1 and Cdk4. Genes indicating S-, G2-, and M-phase are expressed and corresponding columns are indicated in green. All PCR data were normalized to the expression of the housekeeping genes *GAPDH* and *ACTB*. Mean ± SD of three technical replicates are shown.

**Figure S3.**
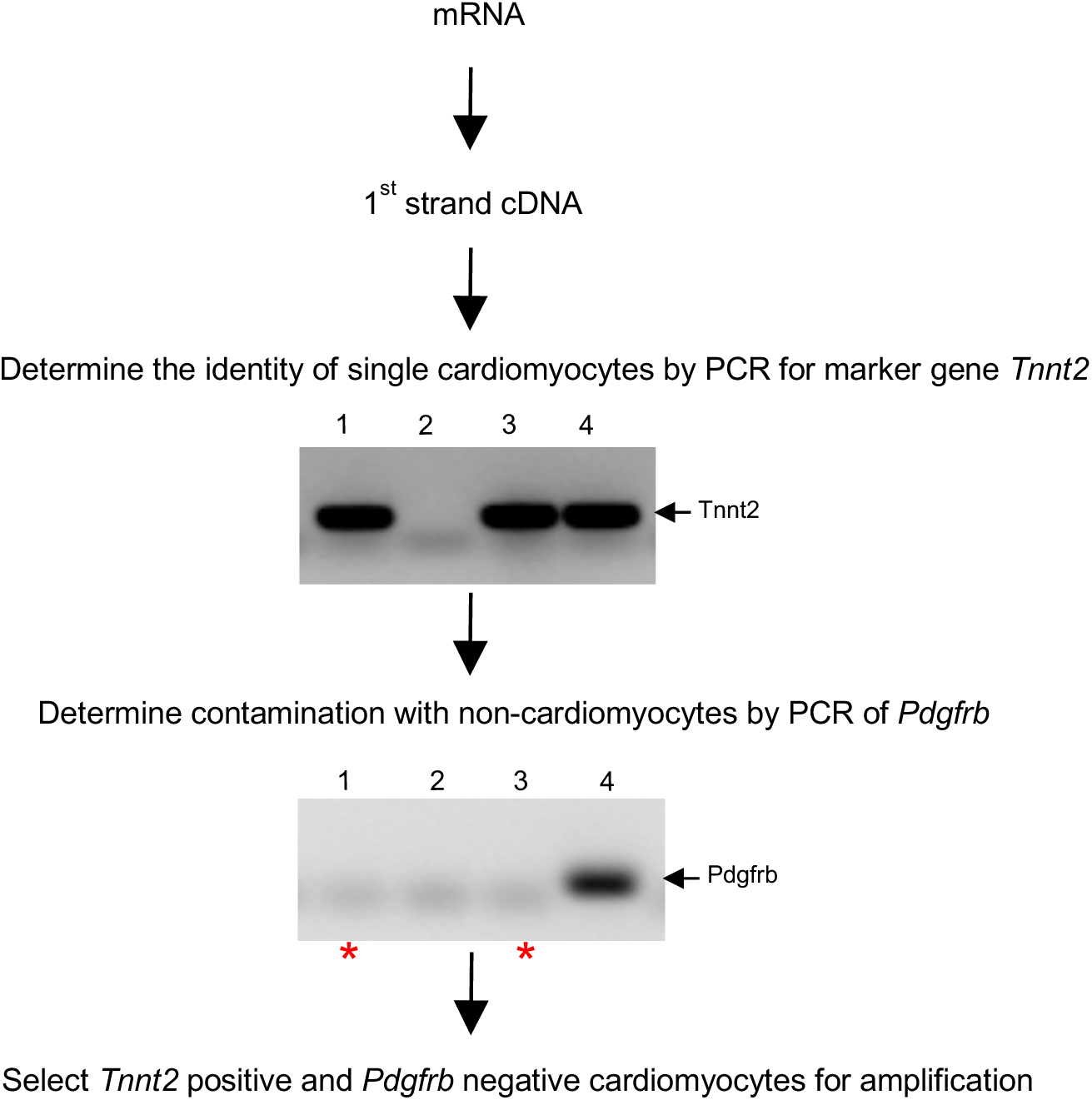
Strategy for identifying single cardiomyocytes. Ventricular cardiac cells were dissociated by enzymatic digestion from Azami green geminin (mAG-Geminin) transgenic mice and sorted by FACS (as explained in **Figure S2**). Following cell lysis, mRNA was reverse transcribed into 1st strand cDNA. Single cell cDNA was subjected to PCR using the cardiomyocyte-specific marker, troponin T2 (*Tnnt2*). Potential contamination with non-cardiomyocytes (i.e. cardiac fibroblasts and endothelial cells) was detected by PCR for platelet derived growth factor receptor β (*Pdgfrb*). Gel electrophoresis of single cells are shown. Single cells showing *Tnnt2* positive and *Pdgfrb* negative were considered as confirmed cardiomyocytes, i.e. cells 1 & 3 (indicated by red asterisks). Cells expressing both genes or neither gene were eliminated. Taken together, our isolation and identification method yielded 40-60% of single cardiomyocytes for single-cell linear amplification and sequencing.

**Fig. S4.**
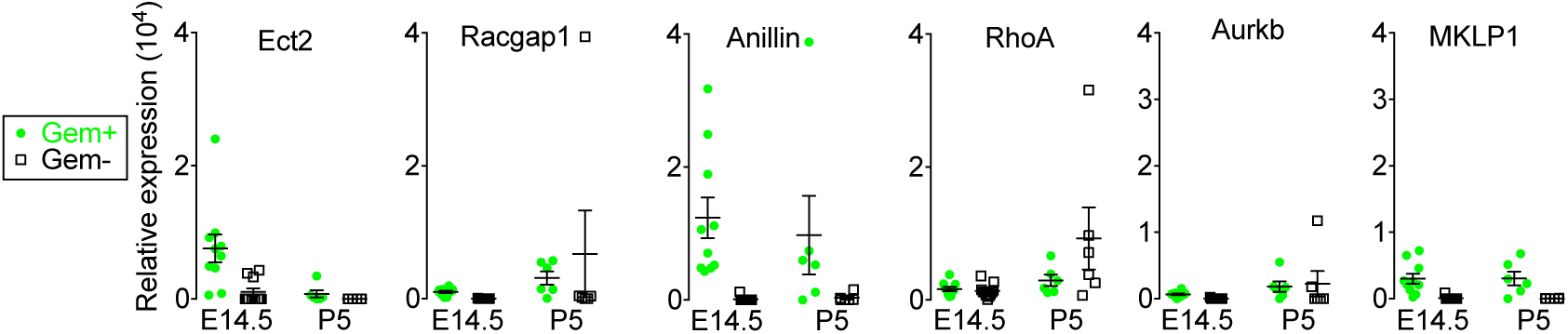
Expression levels of selected cytokinesis genes are graphed for single cardiomyocytes at E14.5 and P5. Mean ± SEM are indicated.

**Figure S5.**
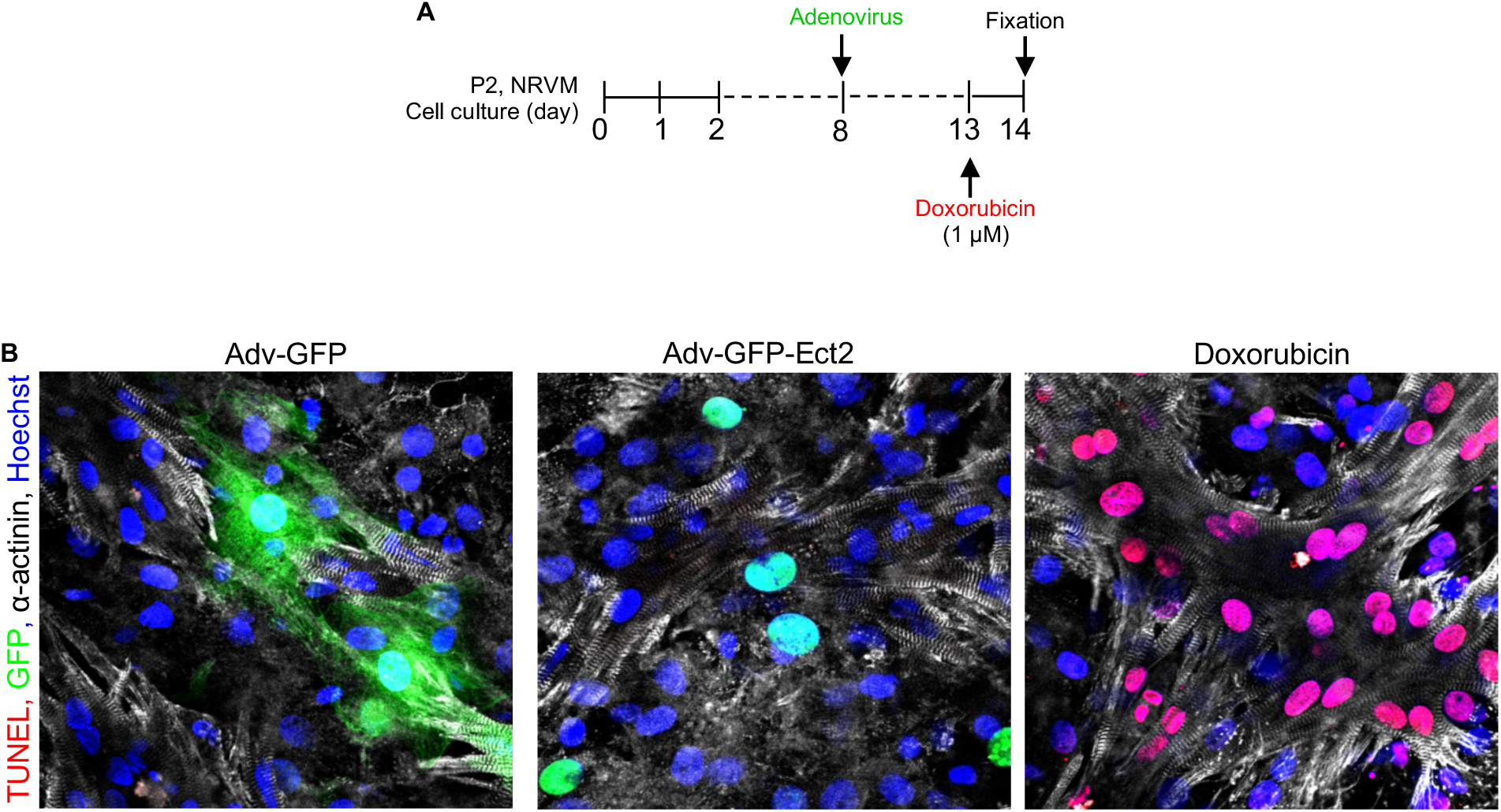
Adenoviral-mediated transduction of Ect2 does not induce apoptosis in cultured neonatal rat ventricular cardiomyocytes (NRVMs). The fresh isolated NRVMs were first cultured for eight days, then Ect2 was expressed in the cells through adenoviral-mediated transduction (Adv-GFP-Ect2, MOI = 200). The transduction of GFP (MOI = 50) was used as negative control. As a positive control, the NRVMs were treated with doxorubicin (1 µM) to induce apoptosis in the cells. (**A**) Design of the apoptosis assay. (**B**) Apoptosis was detected with the ApopTag Red In Situ apoptosis detection kit (EMD Millipore corporation), using Doxorubicin as a positive control.

**Figure S6.**
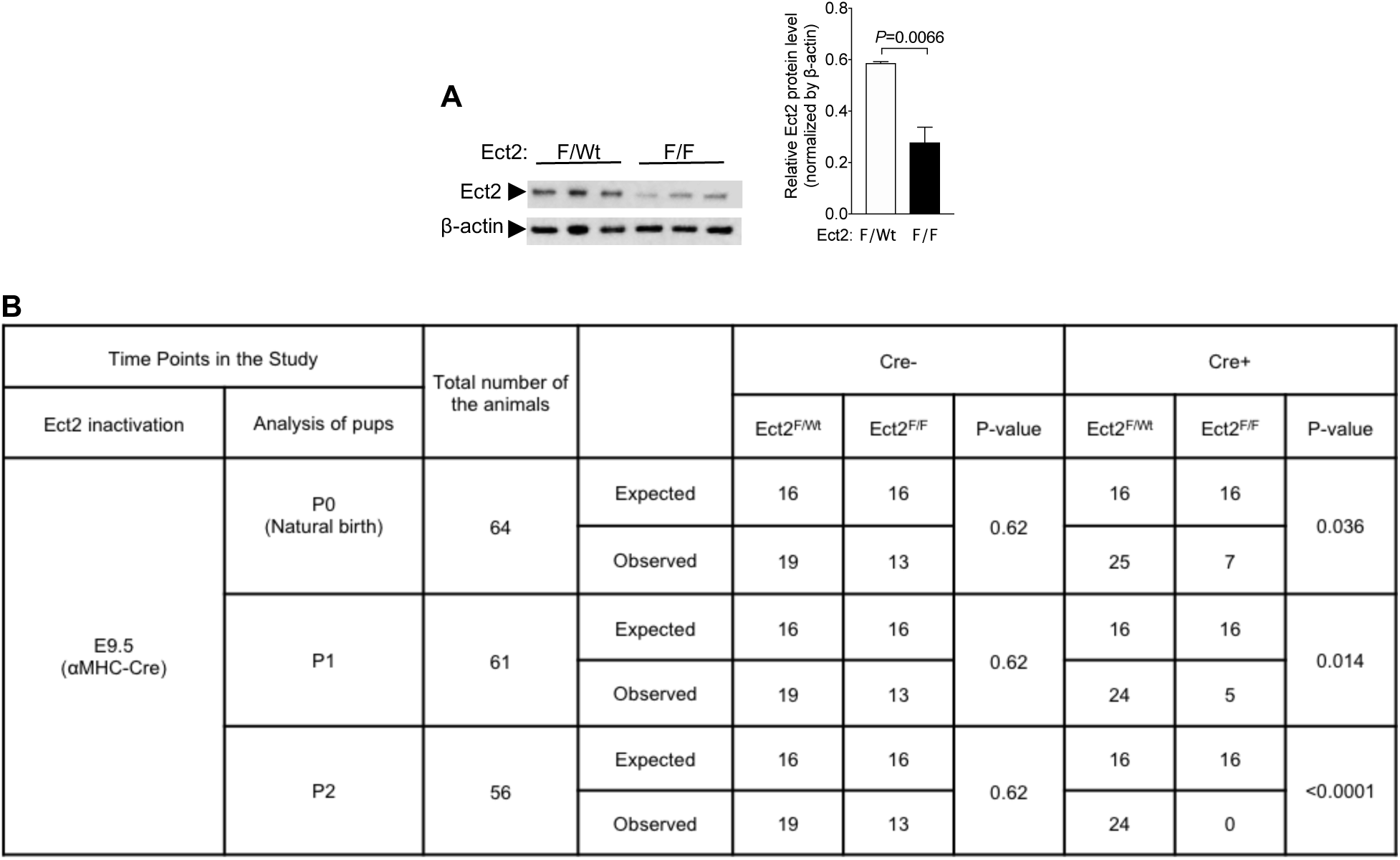
Survival analysis shows that Ect2 gene inactivation in development induces decreased pup viability. To decrease the cardiomyocyte endowment, we inactivated Ect2^Flox^ with αMHC-Cre in embryonic mouse hearts. **(A)** Western blot shows 35% protein reduction in αMHC-Cre; Ect2F/F mice at E16.5 (Ect2F/wt n = 3 hearts, Ect2F/F n = 3 hearts). **(B)** Genotypes and corresponding condition and number of pups recovered are listed. P-values were calculated using Fisher’s exact test. Abbreviations used: F/F, flox/flox; F/Wt, flox/+.

**Figure S7.**
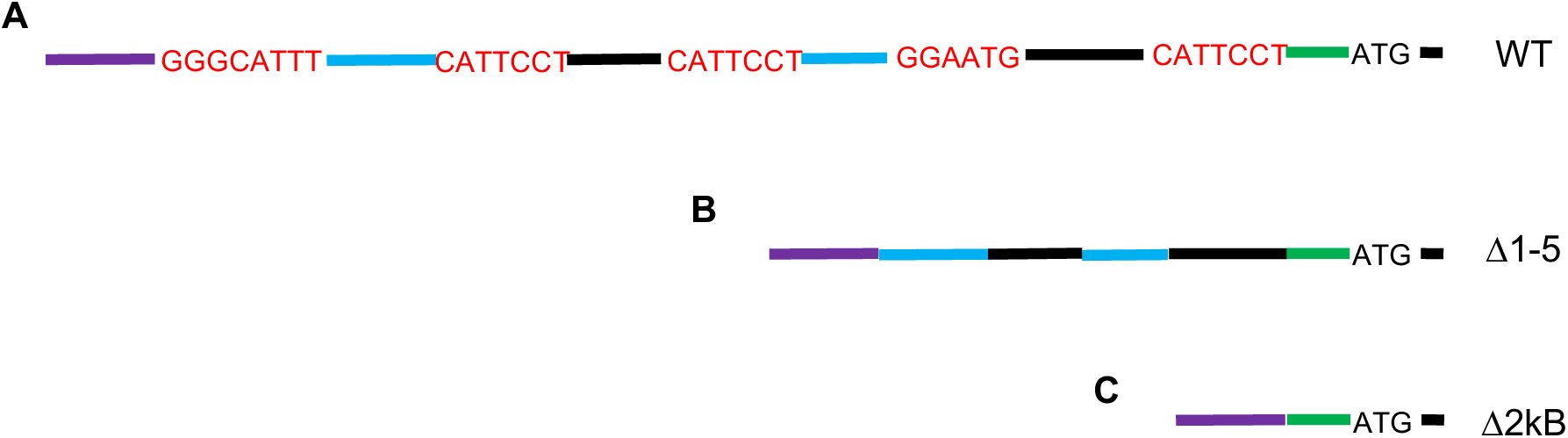
The wild type Ect2 promoter was modified to test the effect of the putative TEAD1/2-binding sites on the Ect2 promoter activity. **(A)** Five putative TEAD1/2-binding sites (red) were detected in the Ect2 promoter region through genome browser. **(B)** All five putative TEAD-binding sites were removed from the Ect2 promoter. **(C)** The continuous 2kB DNA sequence containing all the five TEAD-binding sites was removed.

**Figure S8.**
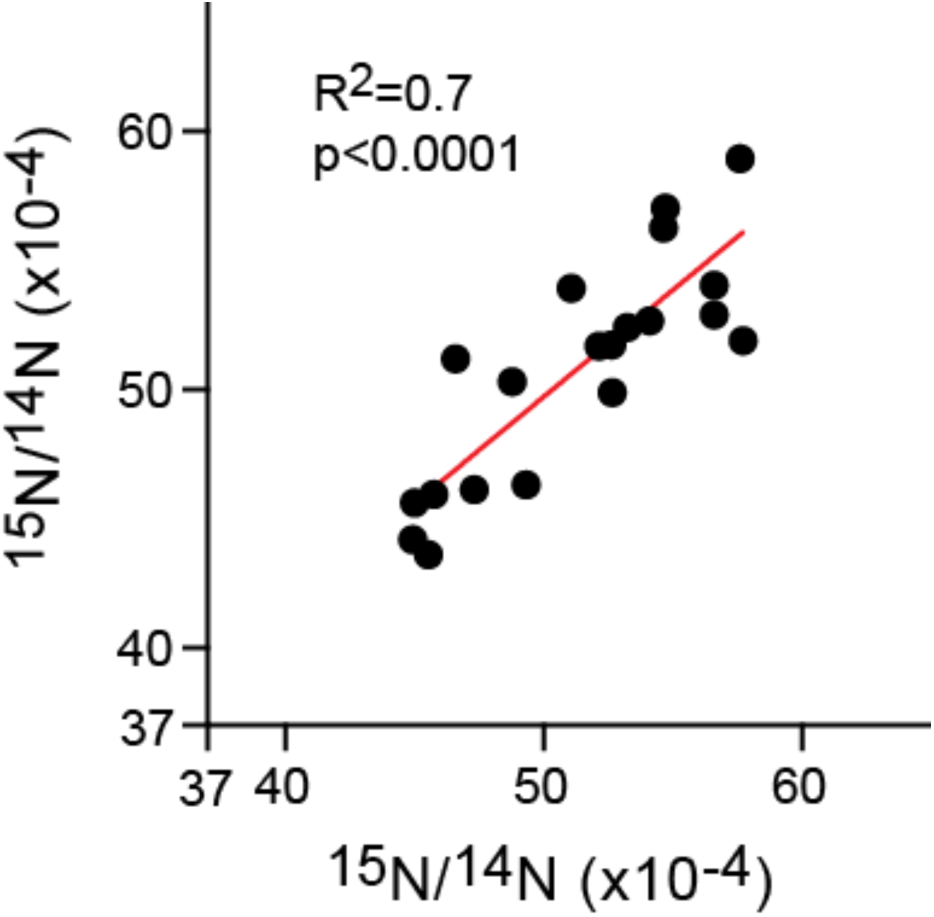
^15^N-thymidine labeling of binucleated cardiomyocytes. The degree of ^15^N-thymidine labeling across all cardiomyocyte nuclei was variable, reflecting in part variable label exposure due to the kinetics of daily label administration. However, within individual nuclei of binucleated cardiomyocytes, the labeling was similar to that shown by the dot plot. Each dot represents a single binucleated cardiomyocyte, with the X/Y coordinates representing the ^15^N/^14^N ratio for each member of the paired nuclei of the binucleated cardiomyocytes. The origin of the graph is at 37 × 10^-4^, which is the natural background ratio of ^15^N relative to the more common stable isotopic variant, ^14^N.

**Figure S9.**
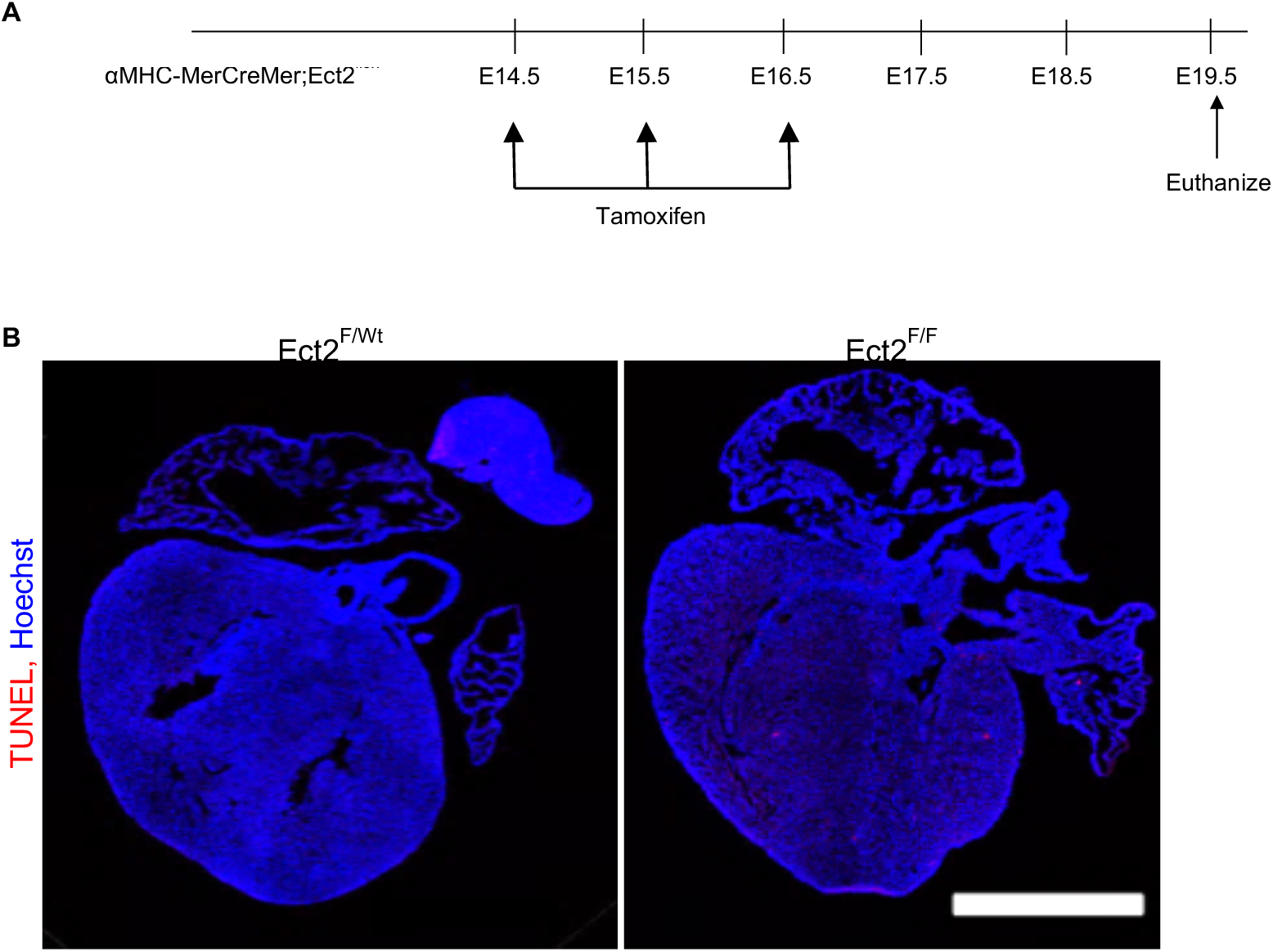
Inactivation of Ect2 gene in fetal cardiomyocytes does not induce apoptosis. **(A)** The Ect2flox gene in the mouse line αMHC-MerCreMer;Ect2^flox^ was inactivated through intraperitoneal (i.p.) injection of tamoxifen (30 μg body weight) to pregnant dams on E14.5, 15.5, and 16.5. Pups were resected and analyzed at E19.5. **(B)** There is no evidence for cardiomyocyte apoptosis detected by TUNEL assay. Scale bar 1 mm.

**Table S1. List of differentially expressed genes encoding 61 different Dbl-homology family Rho-GEF’s between E14.5 and P5 cycling and non-cycling cardiomyocytes.** Ect2 has a p-value of ≤ 0.05 and false discovery rate (FDR) of ≤10%. This table was uploaded separately as an excel file.

**Table S2.**
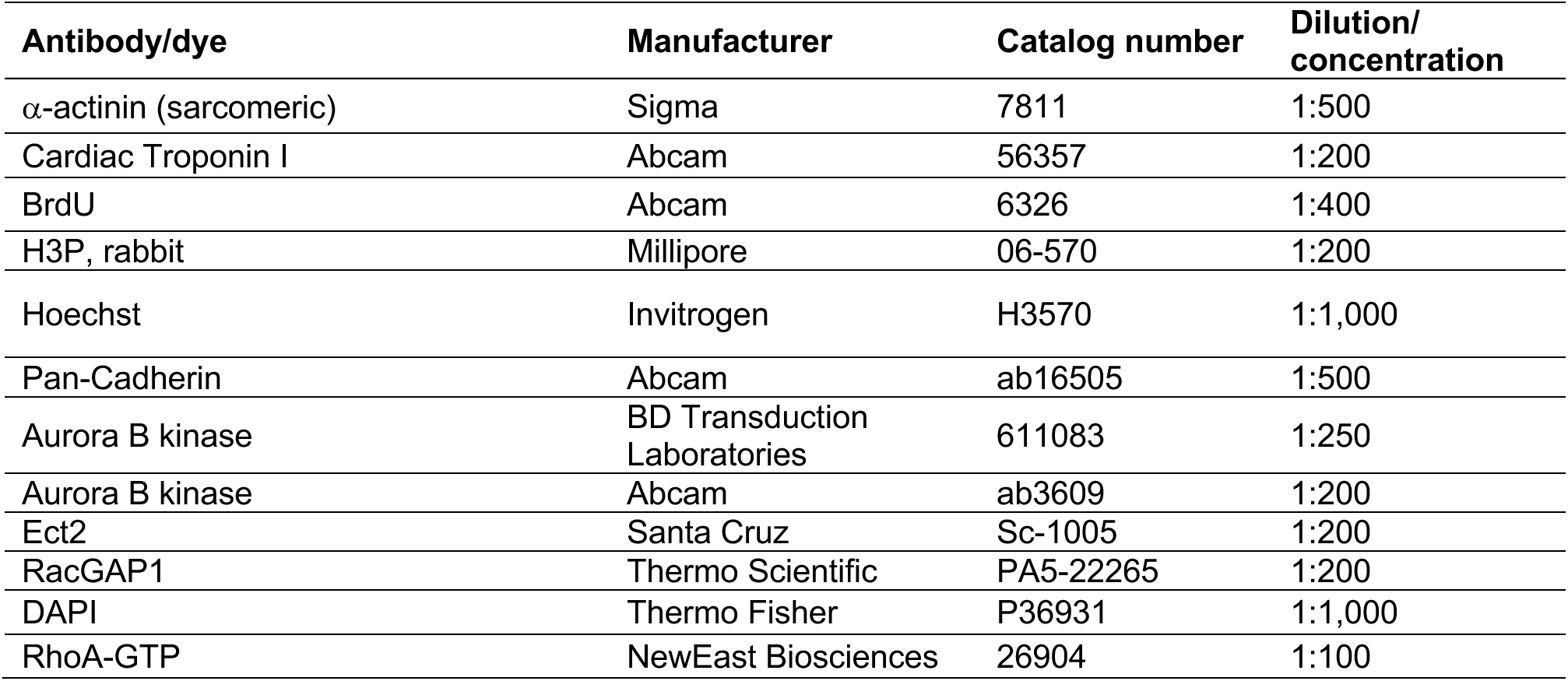
Antibody manufacturers and dilutions

**Table S3.** Image acquisition hardware and settings.

|  | Hardware | Software and Settings |
| --- | --- | --- |
| Fig. 1A | Nikon A1R Confocal 405 nm, 488 nm, 561 nm, 640 nm solid state lasers, 60x Plan Apo Lambda Oil, NA 1.4. | 20–500 msec exposure, Nikon NIS Elements |
| Fig. 1D, E | NanoSIMS 50L (CAMECA) | OpenMIMS version 3.0 |
| Fig. 2A | Olympus IX-81 spinning disk microscope with LUCPLFL $\times 40$, NA 0.6 lens equipped with Hamamatsu EM CCD C9100. | 5–500 msec exposure. Image taken every 20 minutes. |
| Fig. 2C | Nikon A1R Confocal 405 nm, 488 nm, 561 nm, 640 nm solid state lasers, 60x Plan Apo Lambda Oil, NA 1.4. | 20–500 msec exposure, Nikon NIS Elements |
| Fig. 2D | Nikon TiE, Andor Zyla VSC-04468 camera, 60x Plan Apo Lambda Oil, NA 1.4. | 20–500 msec exposure, Nikon NIS Elements |
| Fig. 2E | Nikon A1R Confocal 405 nm, 488 nm, 561 nm, 640 nm solid state lasers, 40x Plan Fluor Oil, NA 1.3. | 20–500 msec exposure, Nikon NIS Elements |
| Fig. 2F | Nikon A1R Confocal 405 nm, 488 nm, 561 nm, 640 nm solid state lasers, 40x Plan Fluor Oil, NA 1.3. | 20–500 msec exposure, Nikon NIS Elements |
| Fig. 2G | Nikon A1R Confocal 405 nm, 488 nm, 561 nm, 640 nm solid state lasers, 40x Plan Fluor Oil, NA 1.3. | 20–500 msec exposure, Nikon NIS Elements |
| Fig. 2H | Nikon A1R Confocal 405 nm, 488 nm, 561 nm, solid state lasers, 60x Plan Apo Lambda Oil, NA 1.4. | 20–500 msec exposure, Nikon NIS Elements |
| Fig. 2K | Nikon TiE, Andor Zyla VSC-04468 camera, Plan Apo $\lambda$ 40x | 20–500 msec exposure, Nikon NIS Elements |
| Fig. 2M | High-resolution ultrasound imaging / micro-injection system (Visual Sonics, Vevo 770) | ImageJ |
| Fig. 2O | Nikon A1R Confocal 405 nm, 488 nm, 561 nm, solid state lasers, 60x Plan Apo Lambda Oil, NA 1.4. | 20–500 msec exposure, Nikon NIS Elements |
| Fig. 2P | Nikon A1R Confocal 405 nm, 488 nm, 561 nm, 640 nm solid state lasers, 40x Plan Fluor Oil, NA 1.3. | 20–500 msec exposure, Nikon NIS Elements |
| Fig. 3C | Nikon A1R Confocal 405 nm, 488 nm, 561 nm, 640 nm solid state lasers, 60x Plan Apo Lambda Oil, NA 1.4. | 20–500 msec exposure, Nikon NIS Elements |
| Fig. 3E | Nikon A1R Confocal 405 nm, 488 nm, 561 nm, 640 nm solid state lasers, 60x Plan Apo Lambda Oil, NA 1.4. | 20–500 msec exposure, Nikon NIS Elements |
| Fig. 3G | Nikon A1R Confocal 405 nm, 488 nm, 561 nm, 640 nm solid state lasers, 60x Plan Apo Lambda Oil, NA 1.4. | 20–500 msec exposure, Nikon NIS Elements |
| Fig. 3H | Nikon A1R Confocal 405 nm, 488 nm, 561 nm, 640 nm solid state lasers, 60x Plan Apo Lambda Oil, NA 1.4. | 20–500 msec exposure, Nikon NIS Elements |
| Fig. 3J | Nikon A1R Confocal 405 nm, 488 nm, 561 nm, 640 nm solid state lasers, 60x Plan Apo Lambda Oil, NA 1.4. | 20–500 msec exposure, Nikon NIS Elements |
| Fig. 3L | Nikon A1R Confocal 405 nm, 488 nm, 561 nm, 640 nm solid state lasers, 60x Plan Apo Lambda Oil, NA 1.4. | 20–500 msec exposure, Nikon NIS Elements |
| Fig. 4C | Bruker, BioSpin 70/30, micro-MRI system | ParaVision 5.1 |
| Fig. 4D | Bruker, BioSpin 70/30, micro-MRI system | ParaVision 5.1 |
| Fig. 4E | Bruker, BioSpin 70/30, micro-MRI system | ParaVision 5.1 |
| Fig. 4F | Bruker, BioSpin 70/30, micro-MRI system | ParaVision 5.1 |
| Fig. 4G | Leica MZ6 with Nikon DS Fi2 | ImageJ |
| Fig. 5C | Nikon A1R Confocal 405 nm, 488 nm, 561 nm, 640 nm solid state lasers, 60x Plan Apo Lambda Oil, NA 1.4. | 20–500 msec exposure, Nikon NIS Elements |
| Fig. 5D | Nikon A1R Confocal 405 nm, 488 nm, 561 nm, 640 nm solid state lasers, 60x Plan Apo Lambda Oil, NA 1.4. | 20–500 msec exposure, Nikon NIS Elements |
| Fig. 5E | Nikon A1R Confocal 405 nm, 488 nm, 561 nm, 640 nm solid state lasers, 60x Plan Apo Lambda Oil, NA 1.4. | 20–500 msec exposure, Nikon NIS Elements |
| Fig. S5 | Nikon A1Rsi Confocal 405 nm, 488 nm, 561 nm, 640 nm solid state lasers, 60x Plan Apo Lambda Oil, NA 1.4. | 20–500 msec exposure, Nikon NIS Elements |
| Fig. S9 | Nikon A1R Confocal 405 nm, 488 nm, 561 nm, 640 nm solid state lasers, 4x Plan Apo VC 20x DIC N2, NA 0.75. | 20–500 msec exposure, Nikon NIS Elements. Large image stitch. |

**Table S4.** Quantification of numeric data.

| Figure | Assay | Number of cardiomyocytes and/or hearts analyzed |
| --- | --- | --- |
| Fig. 1B-C | Quantification of mono/bi/multi nucleated cardiomyocytes in human subjects | ToF/PS patient hearts, n = 12<br>No heart disease hearts, n = 5 |
| Fig. 1F | <sup>15</sup>N-thymidine<sup>+</sup> cardiomyocyte nucleation in ToF/PS patients | ToF/PS patients: 1<br>Mononucleated cardiomyocytes analyzed: 282<br>15N<sup>+</sup> mononucleated cardiomyocytes: 25<br>Binucleated cardiomyocytes analyzed: 104 (208 total nuclei)<br>15N<sup>+</sup> binucleated cardiomyocytes: 20 (40 total nuclei) |
| Fig. 2A | Video microscopy | 52 cardiomyocytes |
| Fig. 2C | Cytokinesis assay:<br>AuroraB//RhoA-GTP colocalization | E14.5: 59 Aurora B kinase positive midbodies<br>P2: 56 Aurora B kinase positive midbodies |
| Fig. 2E | Binucleated BrdU<sup>+</sup> cardiomyocytes | GFP: 407 BrdU<sup>+</sup> cardiomyocytes<br>GFP-Ect2: 285 BrdU<sup>+</sup> cardiomyocytes |
| Fig. 2F | BrdU<sup>+</sup> cardiomyocytes | GFP: 432 cardiomyocytes<br>GFP-Ect2: 443 cardiomyocytes |
| Fig. 2G | H3P<sup>+</sup> nuclei in cardiomyocytes | GFP: 54 H3P<sup>+</sup> nuclei in 2852 cardiomyocyte nuclei<br>Ect2-GFP: 59 H3P<sup>+</sup> nuclei in 3205 cardiomyocyte nuclei |
| Fig. 2H | Percentage of binucleated cardiomyocytes | αMHC-Cre; Ect2<sup>F/Wt</sup> mice: 6<br>αMHC-Cre; Ect2<sup>F/F</sup> mice: 6 |
| Fig. 2I | Percentage of cardiomyocyte nuclei | αMHC-Cre; Ect2<sup>F/Wt</sup> mice: 642 cardiomyocytes<br>αMHC-Cre; Ect2<sup>F/F</sup> mice: 647 cardiomyocytes |
| Fig. 2J | Total number of the cardiomyocytes | αMHC-Cre; Ect2<sup>F/Wt</sup> mice: 12<br>αMHC-Cre; Ect2<sup>F/F</sup> mice: 5 |
| Fig. 2K | Cardiomyocyte size | αMHC-Cre; Ect2<sup>F/Wt</sup> mice: 1138 cardiomyocytes<br>αMHC-Cre; Ect2<sup>F/F</sup> mice: 1015 cardiomyocytes |
| Fig. 2L | Heart-body weight ratio | αMHC-Cre; Ect2<sup>F/Wt</sup> mice: 14<br>αMHC-Cre; Ect2<sup>F/F</sup> mice: 6 |
| Fig. 2M | Ejection fraction | αMHC-Cre-; Ect2<sup>Wt/Wt</sup> mice: 4<br>αMHC-Cre+; Ect2<sup>F/F</sup> mice: 3 |
| Fig. 2N | Pup survival | Fig. S6 |
| Fig. 2O | H3P<sup>+</sup> nuclei in cardiomyocytes | αMHC-Cre; Ect2<sup>F/Wt</sup> mice: 6<br>αMHC-Cre; Ect2<sup>F/F</sup> mice: 6 |
| Fig. 2P | BrdU<sup>+</sup> cardiomyocytes | Wild: 2070 BrdU<sup>-</sup> cells; 84 BrdU<sup>+</sup> cells<br>Ect2KO: 1522 BrdU<sup>-</sup> cells; 70 BrdU<sup>+</sup> cells |
| Fig. 3A | Luciferase assay, Ect2 promoter activity | WT: 3 cultures<br>Δ1-5: 3 cultures<br>Δ2kB: 3 cultures<br>Vector: 3 cultures |
| Fig. 3B | qPCR Ect2 mRNA | Scrambled siRNA: 3 cardiomyocyte isolations<br>TEAD1 siRNA: 4 cardiomyocyte isolations<br>TEAD2 siRNA: 4 cardiomyocyte isolations |
| Fig. 3C | Percentage of binucleated cardiomyocytes | Scrambled siRNA: 3 cardiomyocyte isolations<br>TEAD1 siRNA: 4 cardiomyocyte isolations<br>TEAD2 siRNA: 4 cardiomyocyte isolations |
| Fig. 3D | qPCR Ect2 mRNA | Adv-GFP: 4 cardiomyocyte isolations<br>Adv-YAP1-WT: 4 cardiomyocyte isolations<br>Adv-YAP1-S127A: 4 cardiomyocyte isolations |
| Fig. 3E | Percentage of binucleated cardiomyocytes (BrdU<sup>+</sup>) | Adv-GFP: 4 cardiomyocyte isolations<br>Adv-YAP1-WT: 4 cardiomyocyte isolations<br>Adv-YAP1-S127A: 4 cardiomyocyte isolations |
| Fig. 3F | qPCR Ect2 mRNA | Control: 5 cardiomyocyte isolations<br>Forskolin: 5 cardiomyocyte isolations |
| Fig. 3G | Percentage of multinucleated cardiomyocytes | Control (P4 mice): 6<br>Forskolin (P4 mice): 6<br>Control (P8 mice): 6<br>Forskolin (P8 mice): 6 |
| Fig. 3H | Percentage of multinucleated cardiomyocytes | Wild: 4 hearts<br>DKO: 4 hearts |
| Fig. 3I | Number of cardiomyocytes | Wild: 7 hearts<br>DKO: 5 hearts |
| Fig. 3J | Percentage of H3P<sup>+</sup> cardiomyocytes | Wild: 4 hearts<br>DKO: 4 hearts |
| Fig. 3K | Taqman assay | Wild: 3 hearts<br>DKO: 3 hearts |
| Fig. 3L | Percentage of multinucleated cardiomyocytes | Control (P4 mice): 6<br>Propranolol (P4 mice): 6<br>Control (P8 mice): 6<br>Propranolol (P8 mice): 6 |
| Fig. 3M | Total number of the cardiomyocytes | Control (P4 mice): 7<br>Propranolol (P4 mice): 6<br>Control (P8 mice): 4<br>Propranolol (P8 mice): 4 |
| Fig. 3N | H3P+ nuclei in cardiomyocytes | Control (P4 mice): 6<br>Propranolol (P4 mice): 6 |
| Fig. 3O | Heart-body weight ratio | Control (P4 mice): 7<br>Propranolol (P4 mice): 6<br>Control (P8 mice): 4<br>Propranolol (P8 mice): 4 |
| Fig. 4A | Percentage of Ejection Fraction | Control (PBS): 11<br>Propranolol: 6 |
| Fig. 4C | Percentage of Ejection Fraction | Control (PBS): 5<br>Propranolol: 4 |
| Fig. 4D | Percentage of LV wall thinning caused by adverse remodelling | Control (PBS): 5<br>Propranolol: 4 |
| Fig. 4E | Percentage of LV wall thickening | Control (PBS): 5<br>Propranolol: 4 |
| Fig. 4F | Percentage of Myocardial infarction | Control (PBS): 5<br>Propranolol: 4 |
| Fig. 4G | Scar size | Control (PBS): 5<br>Propranolol: 4 |
| Fig. 4H | Total number of cardiomyocytes | Control (PBS): 5<br>Propranolol: 5 |
| Fig. 4I | Heart-body weight ratio | Control (PBS): 5<br>Propranolol: 6 |
| Fig. 5A | Expression of Ect2 mRNA | ToF/PS: 13 cardiomyocytes<br>Fetal: 71 cardiomyocytes |
| Fig. 5B | Percentage of Ect2 positive cells | ToF/PS: 4 heart samples<br>Fetal: 3 heart samples |
| Fig. 5C | Cardiomyocyte binucleation (BrdU<sup>+</sup>) of cultured human fetal hearts | Control: 4 hearts<br>Forskolin: 4 hearts<br>Dobutamine: 4 hearts<br>Dobutamine+Propranolol: 4 hearts<br>Propranolol: 4 hearts |
| Fig. 5D | Percentage of Ect2+ midbodies | Forskolin: 27 midbodies<br>Dobutamine: 55 midbodies<br>Dobutamine: 26 midbodies |
| Fig. 5E | Cardiomyocyte binucleation (BrdU<sup>+</sup>) of cultured myocardium from ToF/PS patients | Forskolin: 3 hearts<br>Dobutamine: 3 hearts<br>Dobutamine+Propranolol: 3 hearts<br>Propranolol: 3 hearts |

**Table S5.**
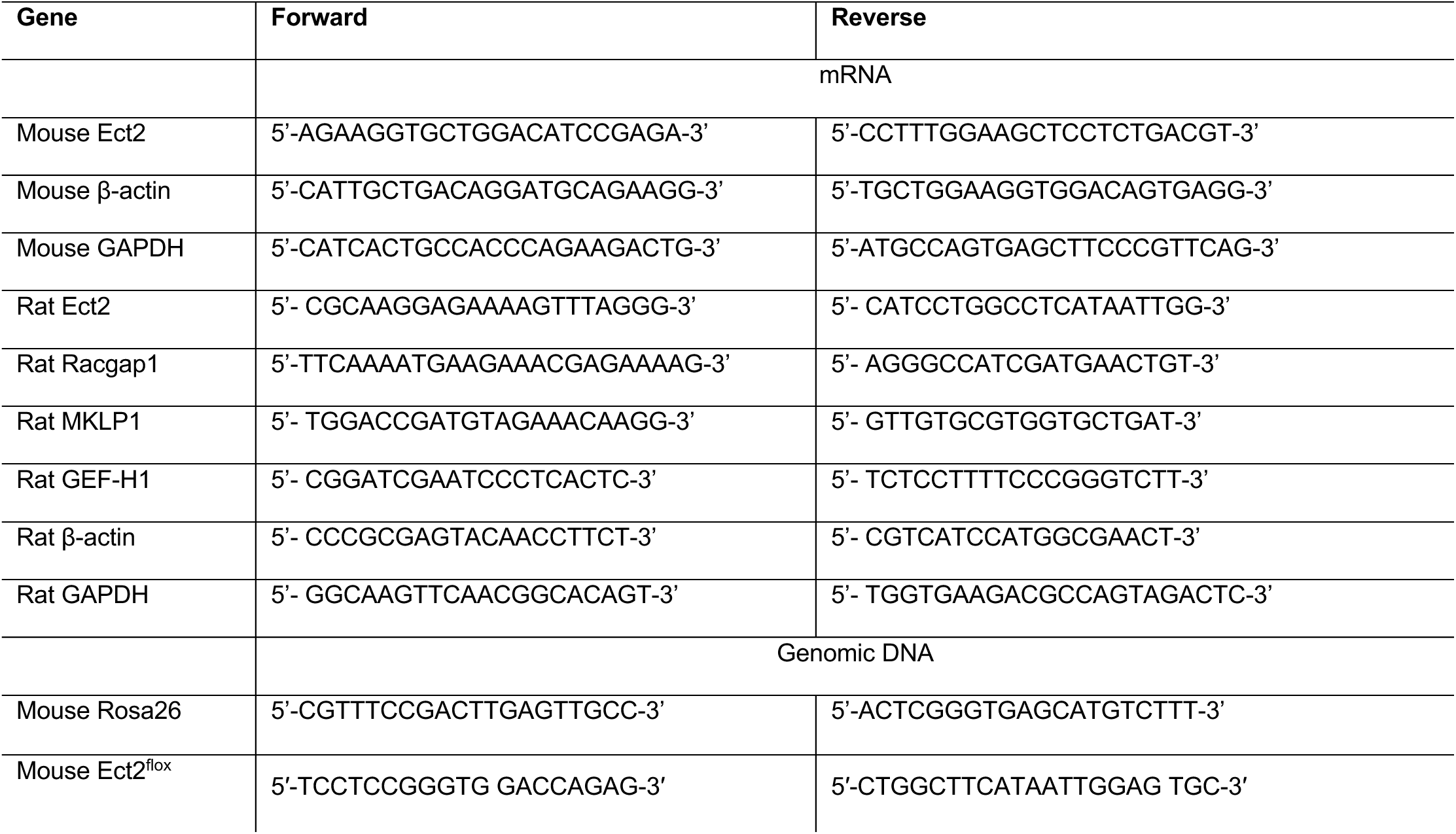
PCR primers and 5’-3’ oligonucleotides

**Table S6.** Animal strains used in this study

| **Gene** | **Species** | **Source** |
| --- | --- | --- |
| FUCCI-red(mKO2-cdt1) | Mouse, C57BL/6 | Miyawaki Lab, RIKEN Institute, Japan |
| FUCCI-green(mAG-Geminin) | Mouse, C57BL/6 | Miyawaki Lab, RIKEN Institute, Japan |
| Ect2<sup>F/F</sup> | Mouse, C57BL/6 | Provided by Dr. Channing Der (University of North Carolina Chapel Hill) |
| αMHC-Cre | Mouse, C57BL/6 | The Jackson Laboratory |
| αMHC-MerCreMer | Mouse, C57BL/6 | The Jackson Laboratory |
| βMHC-YFP(Knock-in) | Mouse, C57BL/6 | Oliver Smithies University of North Carolina Chapel Hill (PNAS) |
| FUCCI-green(mAG-Geminin); βMHC-YFP | Mouse, C57BL/6 | Crossbreeding FUCCI-green(mAG-Geminin) mice with βMHC-YFP mice |
| αMHC-Cre; Ect2<sup>Flox</sup> | Mouse, C57BL/6 | Crossbreeding αMHC-Cre mice with Ect2<sup>Flox</sup> mice |
| αMHC-MerCreMer; Ect2<sup>Flox</sup> | Mouse, C57BL/6 | Crossbreeding αMHC-MerCreMer mice with Ect2<sup>Flox</sup> mice |
| FUCCI-red (mKO2-cdt1; FUCCI-green(mAG-Geminin) | Mouse, C57BL/6 | Crossbreeding FUCCI-red(mKO2-cdt1) mice with FUCCI-green(mAG-Geminin) mice |
| β1-AR<sup>-/-</sup>;β2-AR<sup>-/-</sup> | Mouse, C57BL/6 and (129X1/SvJ x 129S1/Sv)F1-Kitl | The Jackson Laboratory |
| Wild type | Mouse, ICR (CD-1) | Charles River |
| Wild type | Rat, Sprague Dawley | Charles River |

**Table S7.**
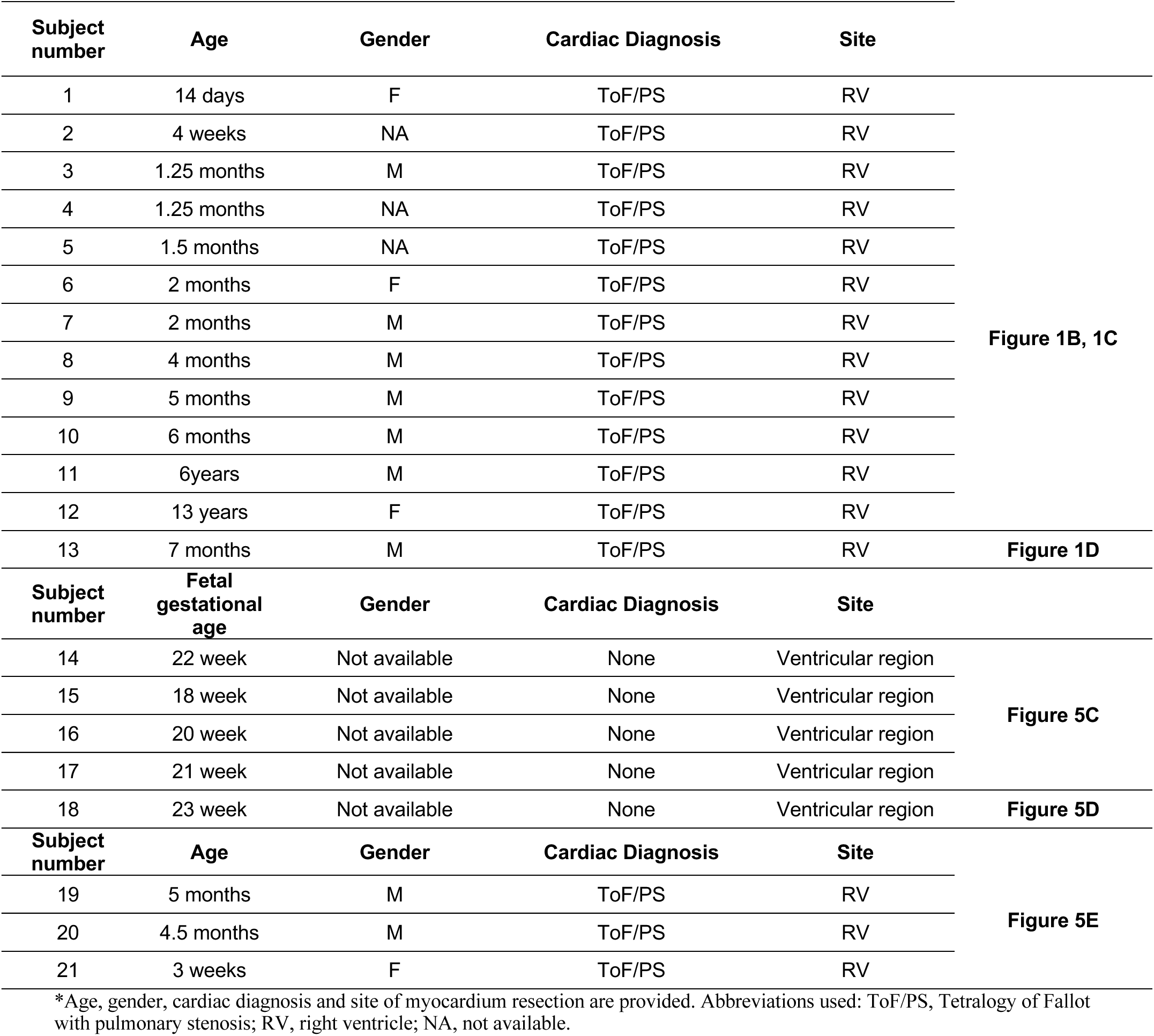
Clinical information corresponding to human samples*

**Table S8.** Ejection fraction of newborn (P0) mice αMHC-Cre; Ect2^flox^ pups. Echocardiography was performed with a Visual Sonics 770. Statistical significance was tested with t test.

| Method | Litter# | Mouse# | Genotype | Ejection fraction (%) | P-value |
| --- | --- | --- | --- | --- | --- |
| Echocardiography | 1 | 1 | αMHC-Cre<sup>-</sup>; Ect2<sup>flox</sup> (Wild) | 90.03 | 0.0068 |
|  |  | 2 |  | 85.41 |  |
|  |  | 3 |  | 85.70 |  |
|  |  | 4 |  | 82.42 |  |
| | | 5 | αMHC-Cre<sup>+</sup>; Ect2<sup>F/F</sup> (Ect2KO) | 66.72 | |
|  |  | 6 |  | 32.39 |  |
|  | 2 | 7 |  | 42.43 |  |

**Video S1. Live cell imaging in NRVM shows that cleavage furrow regression precedes formation of binucleation.** Neonatal rat ventricular cardiomyocytes (NRVM) were imaged by live cell microscopy. Midbody formation is evident between 300-335 minutes, cleavage furrow regression at 355 minutes, and formation of a binucleated cardiomyocyte at 510 minutes.

**Video S2. Live cell imaging of a neonatal cardiomyocyte undergoing division.** Neonatal cardiomyocytes were isolated from mice expressing both elements of the FUCCI reporter in which Cdt1 fused to Orange Fluorescent Protein (Cdt1-mKO2) indicates G1 phase in red, and Geminin fused to Green Fluorescent Protein (mAG-hGem) indicates S-, G2-, and M-phase in green. The video starts with a single cardiomyocyte nucleus in green, indicating that it is in S-, G2-, or M-phase. After division, the cardiomyocyte nucleus turns to red, indicating that it has exited the cell cycle. The last frame is the result of immunofluorescence staining showing BrdU-uptake (blue) and sarcomers stained with α-actinin (red).

**Video S3. Live cell imaging of a neonatal cardiomyocyte undergoing cytokinesis failure.** Neonatal cardiomyocytes were isolated from mice expressing both elements of the FUCCI reporter in which Cdt1 fused to Orange Fluorescent Protein (Cdt1-mKO2) indicates G1 phase in red, and Geminin fused to Green Fluorescent Protein (mAG-hGem) indicates S-, G2-, and M-phase in green. The video starts with a single cardiomyocyte nucleus in green, indicating that it is in S-, G2-, or M-phase. The cell does not divide, both daughter nuclei move together and re-accumulate Cdt1-mKO2, indicating that it has exited the cell cycle. The cells were cultured in the presence of BrdU, whose uptake shows that this cardiomyocyte was in S-phase at the beginning of the video.

**Video S4. Live cell imaging shows appropriate and dynamic localization of GFP-Ect2 during the cell cycle.** NRVM were transduced with adenovirus leading to expression of Ect2-GFP AdV and imaged every 10 minutes. The video shows that GFP-Ect2 is located in the nucleus in interphase (0 minutes), in the cleavage furrow during cytokinesis (60 minutes), in the midbody during abscission (90 minutes), and is associated with completion of cytokinesis (140 minutes).

**Video S5. Ect2 inactivation induces heart failure and death of newborn mice.** Newborn αMHC-Cre^+^;Ect2^flox/flox^ mice (P0) were pale and cool, compared with αMHC-Cre^+^;Ect2^flox/+^ littermates. This mouse (red arrow) died within 48 hours after birth as a result of heart failure.

**Video S6. Heart function of the newborn (P0) wild type mouse pup.** Two-dimensional (2D) B-mode echocardiography recordings covering both ventricles of a newborn (P0) wild type mice shows an ejection fraction of 85.9%.

**Video S7. Heart function of the newborn (P0) mouse pup with Ect2 knockout.** Two-dimensional (2D) B-mode echocardiography recordings covering both ventricles of a newborn (P0) mice with Ect2 knockout shows an ejection fraction of 49.6%.

